# A preoptic circuit triggers rewarming from torpor

**DOI:** 10.64898/2026.04.30.722109

**Authors:** Akinobu Ohba, Madoka Narushima, Chengru Shao, Chi Jung Hung, Hitoshi Uchida, Noriaki Fukatsu, Uyanga Angarag, Sayaka Takemoto-Kimura, Akihiro Yamanaka, Kazuki Tainaka, Daisuke Ono, Yoshifumi Yamaguchi, Hiroaki Wake, Hiroshi Yamaguchi

**Affiliations:** Division of Multicellular Circuit Dynamics, National Institute for Physiological Sciences, Okazaki, Japan; Department of Cell Physiology, Nagoya University Graduate School of Medicine, Nagoya, 466-8550, Japan; Hibernation metabolism, physiology, and development group, Institute of Low Temperature Science, Hokkaido University, Sapporo, 060-0819, Japan; Stress Recognition and Response, Research Institute of Environmental Medicine, Nagoya University, Nagoya, Japan; Department of System Pathology for Neurological Disorders, Brain Research Institute, Niigata University, Niigata, Japan; Department of Anatomy and Molecular Cell Biology, Nagoya University Graduate School of Medicine, Nagoya 466-8550, Japan; Department of Neuroscience, Research Institute of Environmental Medicine, Nagoya University, Nagoya, Japan; Chinese Institute for Brain Research, Beijing (CIBR), Beijing 102206, China; Cold Spring Harbor Laboratory, Cold Spring Harbor, New York, United States

## Abstract

Torpor is an adaptive hypometabolic state that enables homeotherm to survive periods of energetic challenge. This strategy ranges from short bouts of daily torpor to prolonged hibernation. During torpor, animals markedly suppress metabolic rate, body temperature, heart rate, and respiration, while retaining the ability to rewarm. Torpor therefore comprises two critical transitions -entry into a hypometabolic state and active rewarming-both essential for organismal viability. Although neural mechanisms controlling torpor entry have begun to emerge, the circuits that initiate active rewarming and restore euthermia remain poorly defined. Here we identify corticotropin-releasing hormone (Crh)-expressing neurons in the anterodorsal preoptic area (ADP) as a key population for rewarming from fasting-induced torpor in mice. These neurons become active around natural rewarming and are selectively required to limit the depth and duration of torpor. Unlike pathways mediating acute cold defence, stress hyperthermia, or LPS-induced fever, this circuit is dedicated to promoting timely recovery to euthermia. Closed-loop optogenetic activation of ADP^Crh^ neurons during torpor entry rapidly initiates rewarming, and thermographic recordings show that brown adipose tissue (BAT) thermogenesis precedes locomotor arousal. ADP^Crh^ neurons are predominantly GABAergic and project monosynaptically to the lateral preoptic area, whose terminal activation is sufficient to increase body temperature and locomotor activity. Finally, we find robust activation of ADP^Crh^ neurons during rewarming in a hibernator, suggesting conserved logic for exiting deep torpor. Together, our results define a discrete preoptic circuit that drives recovery from torpor and provide a framework for understanding and potentially controlling timely rewarming from profound hypothermia.

## Main

Torpor is a regulated and reversible hypometabolic state that enables homeotherms to survive energetic challenges such as cold and food scarcity. This adaptive strategy is widespread across mammalian and avian lineages and ranges from short daily torpor bouts to prolonged seasonal hibernation^1^. During torpor, animals markedly suppress metabolic rate, body temperature, heart rate, and respiration, while maintaining the ability to rewarm and return to euthermia.

Throughout torpor, the central nervous system operates in a hypometabolic state with reduced electrical and metabolic activity^2,3^. Despite this global suppression, the brain retains the capacity to initiate brown adipose tissue (BAT)-dependent thermogenesis to promote rewarming^4^. While recent studies have begun to define the neural circuits that induce torpor entry^5–9^, the specific neural mechanisms that trigger the thermogenic drive required for active exit from torpor remain largely unknown.

Here, we combine brain-wide activity mapping, a spatial transcriptomic atlas, and causal manipulations to define a hypothalamic circuit that governs torpor rewarming. We identify corticotropin-releasing hormone (*Crh*)-expressing neurons in the anterodorsal preoptic area (ADP) and their projections to the lateral preoptic region as a discrete pathway that limits the depth of torpor and promotes a timely return to euthermia.

## Brain-wide mapping of rewarming-associated regions

To identify the neural circuits involved in rewarming from torpor in mice, we performed a brain-wide mapping of active neurons. Mice were housed at 16 °C and assigned to fed or fasted conditions at the onset of the dark phase. We monitored body surface temperature in real time using an infrared camera. In this screening, torpor bouts were defined as periods in which body surface temperature remained below 28 °C for at least 5 min. All fasted mice entered torpor, and brains were collected approximately 30 to 60 minutes after the onset of rewarming (Fig. 1a, Extended Data Fig. 1a). To minimize circadian effects, we processed fed and fasted mice in pairs and collected the fed control brain at the corresponding time point. The collected brains were subsequently subjected to immunostaining with the anti-phospho S6 (pS6) antibody^10^, an indicator of recent neural activity, and to tissue clearing using iDISCO^11^. We then performed volume imaging with light-sheet microscopy and cell registration using the ClearMap software^12^ (Fig. 1a). To identify brain regions showing differential activity between the two groups, we quantified pS6 signal intensity and calculated fold induction and statistical significance for each brain region. The resulting volcano plot highlights several brain regions that were robustly activated during rewarming, with the anterodorsal preoptic area (ADP) appearing as one of the most significantly enriched regions (Fig. 1b). Consistent with our automated whole-brain analysis (Fig. 1c, Extended Data Fig. 1b), histological validation confirmed a dense accumulation of pS6-positive neurons specifically in the ADP of rewarming mice, whereas minimal activity was observed in control mice (Fig. 1d). Next, to pinpoint the molecular identity of these rewarming-activated neurons in the ADP, we leveraged a spatial transcriptomic cell atlas of the mouse preoptic area^13^. From the atlas, we extracted neuronal clusters spatially enriched around the ADP and compared their predicted distributions with the pS6 map. Inhibitory clusters I-4 and I-10, and the excitatory cluster E-6 showed the closest match to the distribution of rewarming-associated pS6-positive cells in the ADP (Fig. 1e, Extended Data Fig. 1c). We then performed differential expression analysis for each candidate cluster versus all other preoptic clusters, identifying *Nos1* as a common marker for E-6 and I-4, and *Crh* as a leading marker for I-10. Immunostaining further showed that both *Nos1*-positive and *Crh*-positive cells were distributed in and around the ADP (Fig. 1f). To determine the molecular identity of rewarming-activated neurons in vivo, we next generated Crh-iCre; Ai14^14^ reporter mice to genetically label *Crh*-positive neurons with tdTomato. We first confirmed that *Crh*-positive neurons are largely distinct from *Nos1*-positive neurons within the ADP, with only limited overlap between the two populations (Extended Data Fig. 1d). Following torpor rewarming, we found that 41.5% of pS6-positive cells co-expressed tdTomato. In contrast, Nos1 immunoreactivity was detected in only 6.2% of the activated neurons (Fig. 1g, h). Consistent with this, a significant fraction (34.0 ± 4.7%) of *Crh*-positive neurons were positive for pS6 during rewarming, whereas only a small proportion (5.8 ± 0.9%) of *Nos1*-positive neurons were activated (Extended Data Fig. 1e). These results indicate that *Crh*-positive neurons constitute a major component of the ADP population recruited during torpor rewarming.

**Fig. 1:**
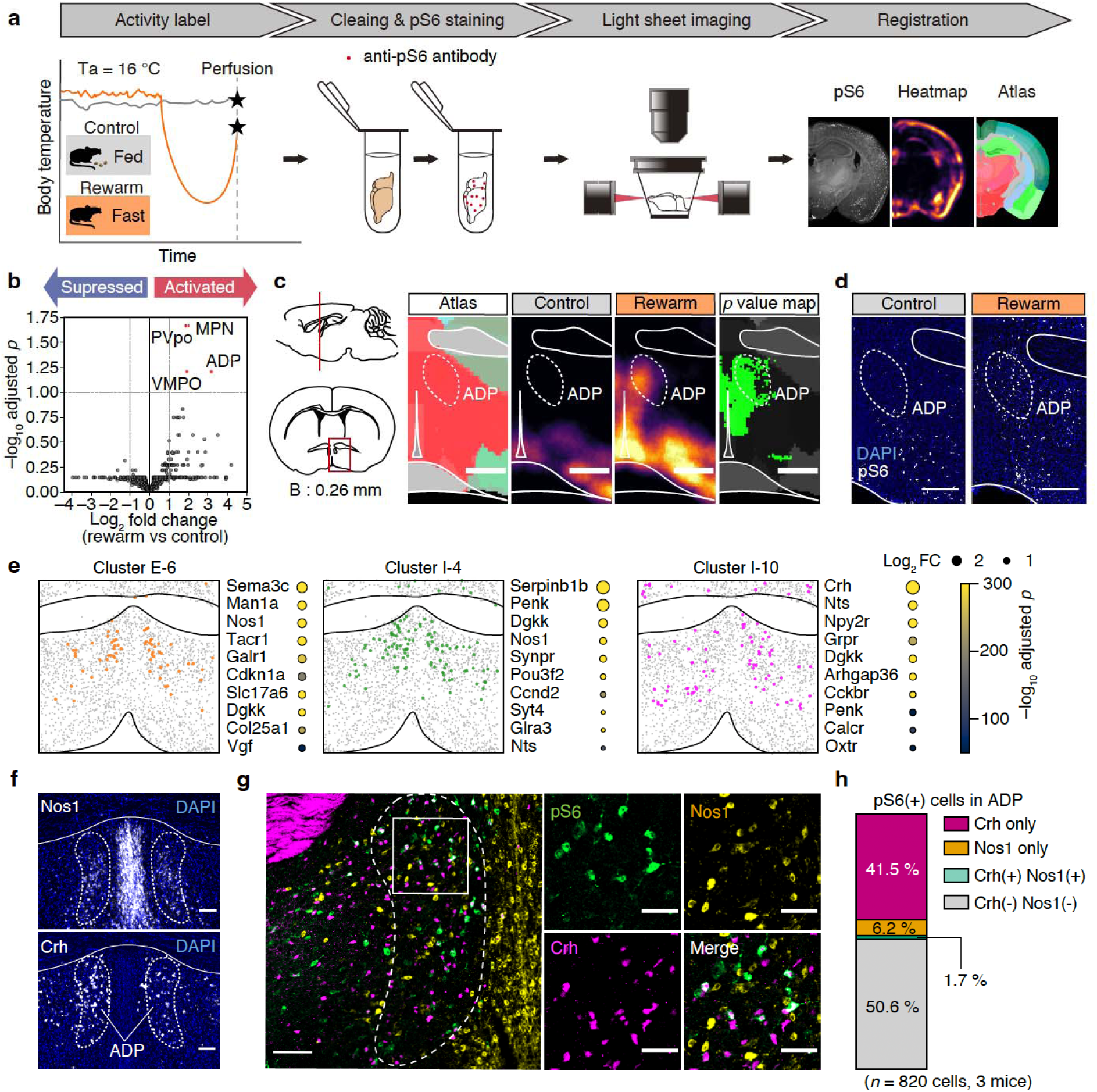
ADPCrh neurons are activated during rewarming. **a,** Experimental workflow for brain-wide activity mapping associated with rewarming. C57BL/6J female mice were transferred to fresh cages at ZT12 and maintained at an ambient temperature of 16 °C under either fed (control, *n* = 7) or fasted (rewarm, *n* = 7) conditions. Body surface temperature was monitored in real time using an infrared camera, and brains were collected approximately 30–60 min after the onset of rewarming (see Methods). Brains were then immunostained for phospho-S6 (pS6), optically cleared, imaged by light-sheet fluorescence microscopy, and registered to the Allen Brain Atlas. **b**, Volcano plot showing brain-wide changes in pS6 signal density in rewarm mice relative to controls. Group comparisons were performed using two-sided Welch’s t-tests followed by Benjamini-Hochberg false discovery rate correction. Each dot represents one brain region. Significant regions are highlighted in red, and non-significant regions are shown in grey. Highlighted regions include ADP (anterodorsal preoptic nucleus), PVpo (periventricular hypothalamic nucleus, preoptic part), VMPO (ventromedial preoptic nucleus), and MPN (medial preoptic nucleus). **c**, Schematic illustration at bregma + 0.26 mm showing the ADP region (left), mean pS6 density heatmaps for control and rewarm conditions (middle), and the voxel-wise *p*-value map (right). For p-value map generation, voxel-wise group differences were assessed by t-test, and voxels with *P* < 0.001 are shown for visualization. Green indicates voxels showing significantly higher activity during rewarming than in the control condition. Scale bar, 500 µm. **d**, Representative fluorescence images showing pS6-positive cells around the ADP in control (left) and rewarm (right) conditions. Scale bar, 300 µm. **e**, Spatial maps of ADP-enriched clusters from a published spatial transcriptomic atlas (Dryad, doi:10.5061/dryad.8t8s248), corresponding to Moffitt et al^13^. Among the seven ADP-enriched clusters, the three clusters (E-6, I-4, and I-10) were selected on the basis of spatial similarity to the pattern of pS6-positive cells in **d**. For each cluster, enriched genes relative to all other neuronal clusters are listed. Bubble size is proportional to the positive log2 fold change, and bubble colour indicates –log10 (adjusted *P* value) based on the Mann–Whitney U test. *P* values were corrected for multiple comparisons using the Benjamini–Hochberg false discovery rate procedure. **f**, Representative images showing the distribution of Nos1 and Crh immunoreactivity in the ADP of wild-type mice. Scale bars, 100 µm. **g**, Representative images showing overlap among pS6, Nos1 and Crh-positive neurons in the ADP. Brains from Crh-iCre; Ai14 reporter mice were collected after rewarming and immunostained for pS6 and Nos1. Scale bars, 100 µm (left) and 50 µm (right). **h**, Proportion of Crh- and Nos1-expressing cells among pS6-positive cells in the ADP (*n* = 820 cells from 3 mice).

## ADP^Crh^ neurons promote timely rewarming

To dissect the functional contributions of the two candidate populations, we employed chemogenetic inhibition. We first targeted ADP neurons expressing *Nos1*(ADP^Nos1^) by injecting a Cre-dependent AAV encoding the Gi-coupled DREADD hM4Di into the ADP of Nos1-cre mice. Inhibiting ADP^Nos1^ neurons produced no detectable changes in torpor depth and bout duration (Extended Data Fig. 2). We therefore focused on ADP neurons expressing *Crh* (ADP^Crh^) in subsequent loss-of-function experiments. To inhibit ADP^Crh^ neurons, we injected a Cre-dependent AAV encoding hM4Di-mCherry into the ADP of Crh-iCre mice (Fig. 2a, b). After intraperitoneal implantation of a temperature logger, mice were fasted at dark onset and injected with clozapine *N*-oxide (CNO) 3.5 h after food removal (Fig. 2c). In Cre-dependent mCherry-only controls, CNO did not alter torpor, indicating that CNO itself does not account for the phenotype (Fig. 2c). In contrast, inhibiting ADP^Crh^ neurons prolonged and deepened the torpor bout (Fig. 2c-e, Extended Data Fig. 3a). Consistently, taCasp3-mediated ablation^15^ of ADP^Crh^ neurons produced longer and deeper torpor bouts compared with controls (Fig. 2f-j, Extended Data Fig. 3b). Together, these results indicate that ADP^Crh^ neurons are required to limit the depth and duration of torpor and to promote timely rewarming.

**Fig. 2:**
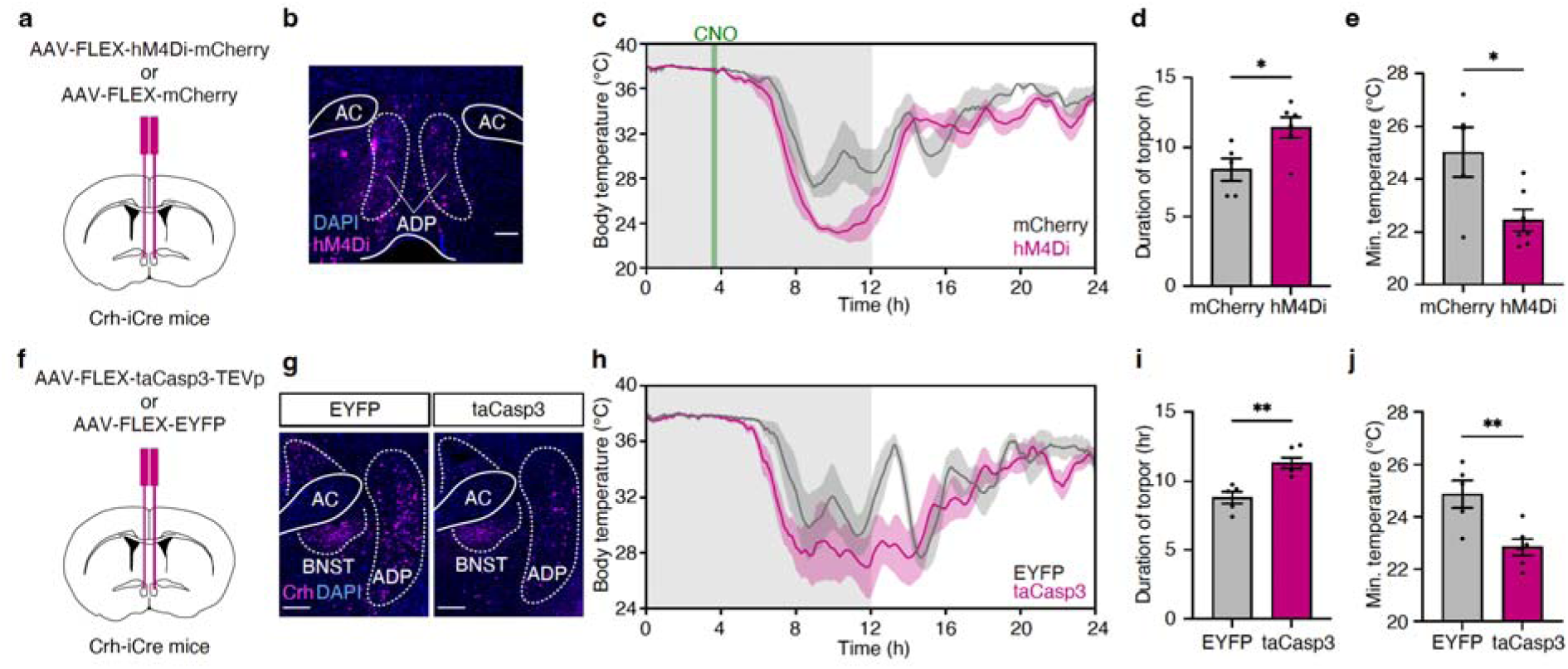
Necessity of ADP^Crh^ neurons for timely rewarming from torpor. **a**, Experimental schematic for chemogenetic inhibition. **a**, Cre-dependent AAV encoding hM4Di-mCherry (AAV9-CAG-FLEX-hM4Di-mCherry) or mCherry control (AAV9-CAG-FLEX-mCherry) was bilaterally injected into the ADP of Crh-iCre mice. **b**, Representative fluorescence image showing viral expression in the ADP. AC, anterior commissure. Scale bar, 200 µm. **c**, Body temperature during 24-hour fasting with CNO administration. Fasting was initiated at ZT12, and CNO was injected 3.5 h after fasting onset (green line). Grey shading indicates the dark phase. Traces show mean ± s.e.m. **d**, Torpor duration, defined as the total time spent at body temperature < 33 °C during the 24-h fasting period (mCherry, *n* = 5 mice; hM4Di, *n* = 7 mice). **e**, Minimum body temperature during the 24-hour fasting. **f**, Experimental schematic for neural ablation. Cre-dependent AAV encoding taCasp3-TEVp (AAV2-FLEX-taCasp3-TEVp) or EYFP control (AAV2-EF1α-DIO-EYFP) were bilaterally injected into the ADP of Crh-iCre mice. **g**, Representative fluorescence images showing Crh immunofluorescence around the ADP (EYFP control, left; taCasp3, right). Scale bar, 200 µm. **h**, Body temperature during 24-h fasting. Grey shading indicates the dark phase. Traces show mean ± s.e.m. **i**, Torpor duration (EYFP, *n* = 5 mice; taCasp3, *n* = 6 mice). **j**, Minimum body temperature. Bars indicate mean ± s.e.m.; each dot represents one mouse. Statistical significance was assessed using two-sided unpaired t-tests. \**P* < 0.05, \*\**P* < 0.01.

To determine whether ADP^Crh^ neurons function as general thermoregulatory effectors or are specifically involved in torpor rewarming, ADP^Crh^ neuron-ablated mice were exposed to various thermogenic stimuli. The ablation of ADP^Crh^ neurons did not impair the baseline body temperature (Extended Data Fig. 4a) nor the maintenance of body temperature during acute cold exposure at 4 °C for 12 hours (Extended Data Fig. 4b). Additionally, both cage change stress-induced hyperthermia and the fever response to lipopolysaccharide were indistinguishable between ablated and control mice (Extended Data Fig. 4c, d). These results indicate that ADP^Crh^ neurons are not required for cold defence, stress responses, or fever, but instead play a specific role in mediating torpor rewarming.

## ADP^Crh^ neurons are sufficient for rewarming

Having established the necessity of ADP^Crh^ neurons for natural rewarming, we next examined whether their activation is sufficient to drive rewarming from torpor. We bilaterally injected a Cre-dependent AAV encoding ChRmine^16^ into the ADP of Crh-iCre mice and implanted optical fibres above the injection site (Fig. 3a, b). Because the timing of torpor entry varies across individual animals, we used a closed-loop system to deliver photostimulation with high temporal precision. Light was delivered continuously whenever maximum body surface temperature remained below 33 °C (Fig. 3b, c). In Cre-dependent hrGFP controls, light delivery did not alter the torpor–arousal cycle, indicating that photostimulation alone does not affect torpor phenotype (Fig. 3d). In contrast, closed-loop activation of ADP^Crh^ neurons rapidly initiated rewarming, increasing the nadir temperature (Fig. 3d–f). Post hoc histology showed robust c-Fos expression in ADP^Crh^ neurons, confirming the specific activation of the targeted population (Fig. 3b, Extended Data Fig. 5). Taken together, these results indicate that activation of ADP^Crh^ neurons is sufficient to trigger efficient rewarming from torpor. To investigate the effector mechanism engaged by ADP^Crh^ neurons, we stimulated this population under euthermic conditions. Optogenetic activation of ADP^Crh^ neurons increased body surface temperature by 1.24 ± 0.30 °C and increased locomotor activity (Fig. 3g-k). To determine whether rewarming is driven by brown adipose tissue (BAT)-dependent thermogenesis or is secondary to locomotor activity, we next performed thermographic recordings at 1 Hz during optogenetic activation of ADP^Crh^ neurons. Thermal imaging revealed a rapid increase in temperature in the interscapular region, the anatomical site of BAT (Fig. 3l, m). Crucially, quantitative analysis demonstrated that the increase in BAT temperature significantly preceded the onset of locomotor activity (Fig. 3n, o). This temporal dissociation indicates that ADP^Crh^ neurons trigger rewarming primarily via BAT-dependent thermogenesis, rather than as a consequence of behavioural arousal.

**Fig. 3:**
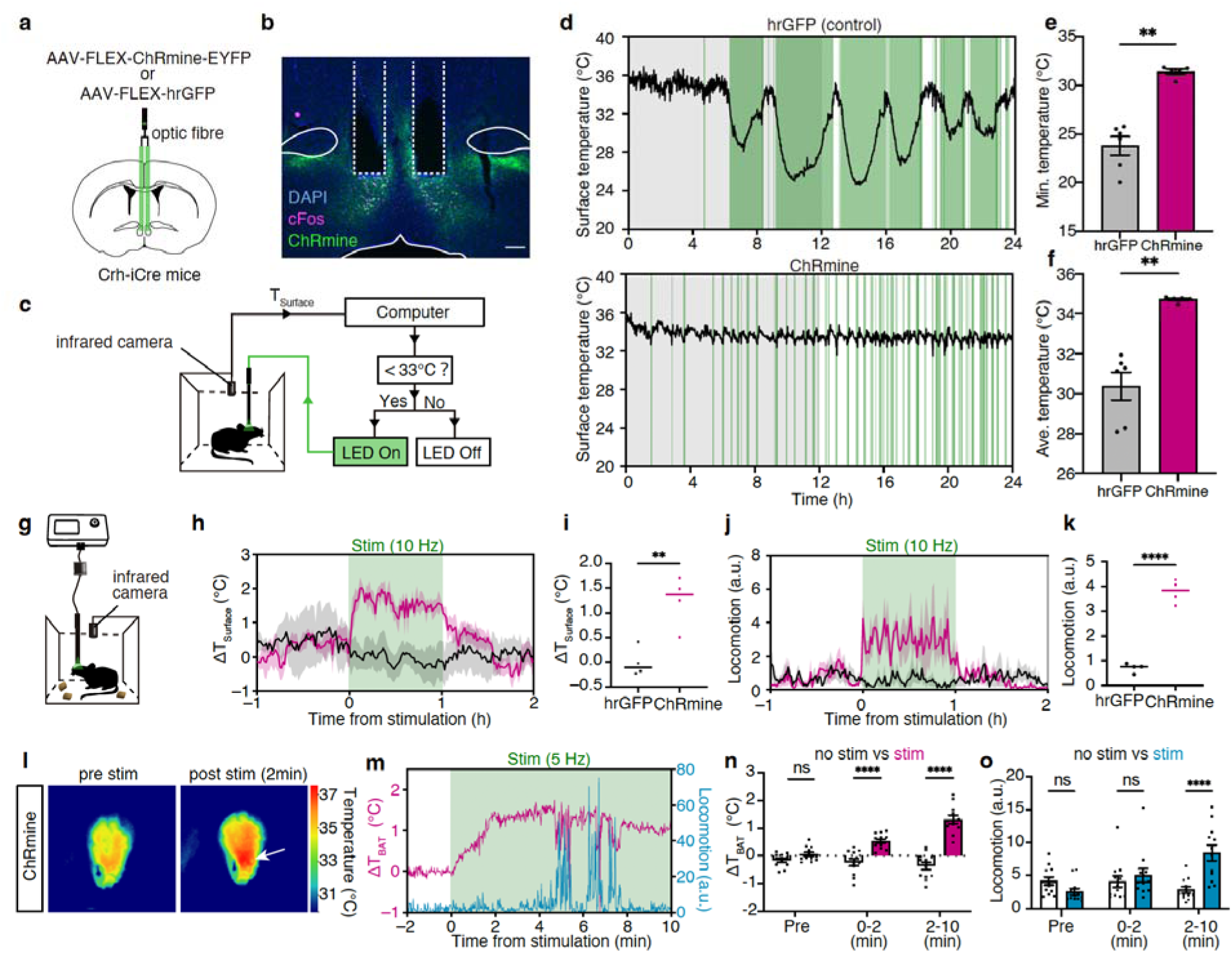
Activation of ADP^Crh^ neurons promotes rewarming from torpor. **a**, Experimental schematic for optogenetic activation. **a,** Cre-dependent AAV encoding ChRmine-EYFP (AAV9-CAG-FLEX-ChRmine-EYFP) or hrGFP control (AAV-CMV-FLEX-hrGFP) was bilaterally injected into the ADP of Crh-iCre mice, and optic fibre was implanted above the ADP. **b**, Representative fluorescence image showing viral expression in the ADP and fibre trace. Scale bar, 200 µm. **c**, Closed-loop optogenetic stimulation paradigm during fasting. Surface temperature was monitored in real time by an infrared camera, and stimulation was triggered when the temperature fell below 33 °C. **d**, Representative surface temperature traces during fasting in hrGFP control (top) and ChRmine (bottom) mice. Green shading indicates epochs of light delivery. Grey shading indicates the dark phase. **e**, Minimum surface temperature during the 24-hour fasting (hrGFP, *n* = 6 mice, ChRmine, *n* = 5 mice, two-sided Mann–Whitney *U* test). **f**, Mean surface temperature during the photostimulation period. (hrGFP, n = 6 mice; ChRmine, n = 5 mice; two-sided Mann–Whitney *U* test). **g**, Schematic of the experimental setup for optogenetic stimulation under euthermic, ad libitum-fed conditions. Surface temperature was recorded using an infrared camera. **h**, ΔT_Surface_ trace during 10 Hz stimulation. Trace show mean ± s.e.m. **i**, Mean ΔT_Surface_ during the 1-hour stimulation window (hrGFP, n = 4 mice, ChRmine, n = 4 mice). For ΔT_Surface_ quantification, baseline temperature was defined as the mean body surface temperature during the 1-h period between ZT0 and ZT1. **j**, Locomotion trace during 10 Hz stimulation. Trace show mean ± s.e.m. **k**, Mean locomotion during the 1-hour stimulation window (hrGFP, *n* = 4 mice; ChRmine, *n* = 4 mice; two-sided unpaired *t*-test). **l**, Representative infrared images before and 2-min after 5 Hz stimulation. The arrow indicates the interscapular BAT region showing an increase in temperature after stimulation. **m**, Representative traces of ΔT_BAT_ (magenta) and locomotion (blue) aligned to the onset of 5 Hz photostimulation (green shading). **n,** ΔT_BAT_ quantified in epochs relative to stimulation onset (Pre, 0–2 min, and 2–10 min) (*n* = 12 trials from 3 mice). For ΔT_BAT_ quantification, baseline temperature was defined as the mean temperature during the 5-min period from 10 to 5 min before the onset of each stimulation epoch. The no-stimulation period was defined as the 10-min window beginning 20 min after the end of each stimulation epoch. **o**, Locomotor activity quantified in the same epochs as in **n**. P values were calculated using two-sided Mann–Whitney U tests for each time window (Pre, 0–2 min, and 2–10 min), followed by Bonferroni correction for three comparisons. Bars indicate mean ± s.e.m.; each dot represents one mouse in **e,f,i,k** and one trial in **n,o**. \*\**P* < 0.01, \*\*\*\**P* < 0.0001; ns, not significant.

We next examined whether ADP^Crh^ neurons are naturally recruited around torpor rewarming. To monitor the temporal dynamics of ADP^Crh^ neurons using fibre photometry calcium imaging, we unilaterally injected a Cre-dependent AAV encoding jGCaMP8s into the ADP of Crh-iCre mice and implanted an optical fibre above the injection site (Extended Data Fig. 6a-d). To avoid confounding effects of temperature-dependent changes in the GCaMP signal, we analysed calcium transient events using a local z-score calculated within a 5-min sliding window and defined events as those exceeding 3 standard deviations (>3 s.d.). We then quantified event magnitude as the area under the curve (AUC). We found that event magnitude was greater during rewarming than during the torpor-entry phase (Extended Data Fig. 6e, f), indicating increased ADP^Crh^ neuronal activity during this period. These data are consistent with the activation of ADP^Crh^ neurons during torpor rewarming.

## The ADP→LPO pathway mediates the increase in body temperature and locomotor activity

We next sought to identify the downstream pathway mediating the thermogenic response evoked by activation of ADP^Crh^ neurons. To delineate the output architecture of ADP^Crh^ neurons, we labelled them with a Cre-dependent EYFP and performed whole-brain clearing. Light-sheet fluorescence imaging revealed widespread projections from the ADP to multiple hypothalamic and brainstem nuclei (Fig. 4a). We then expressed synaptophysin-EGFP in ADP^Crh^ neurons to map putative presynaptic terminals with higher resolution (Fig. 4b). Fluorescence imaging revealed dense terminal innervation in several candidate thermoregulatory regions, including the lateral and dorsomedial hypothalamus (LH/DMH), supramammillary nucleus (SuM), and the lateral preoptic area (LPO) (Fig. 4b). We next asked which projection is sufficient to induce the thermogenic response. We expressed ChRmine in ADP^Crh^ neurons and delivered bilateral photostimulation to axon terminals in candidate target regions with dense terminal fields, such as the LPO, LH/DMH, and SuM (Fig. 4c). Stimulation of ADP^Crh^ terminals in the LPO robustly increased body temperature and locomotor activity (Fig. 4d). In contrast, terminal stimulation in the LH/DMH or SuM produced little or no change in body temperature and locomotor activity (Fig. 4e, f). Fibre placements were verified histologically (Extended Data Fig. 7). These results identify the ADP^Crh^→LPO pathway as a key downstream route sufficient to drive the thermogenic and behavioural effects of ADP^Crh^ neuron activation. Consistently, closed-loop activation of the ADP^Crh^→LPO pathway during torpor rapidly triggered rewarming (Extended Data Fig. 8).

**Fig. 4:**
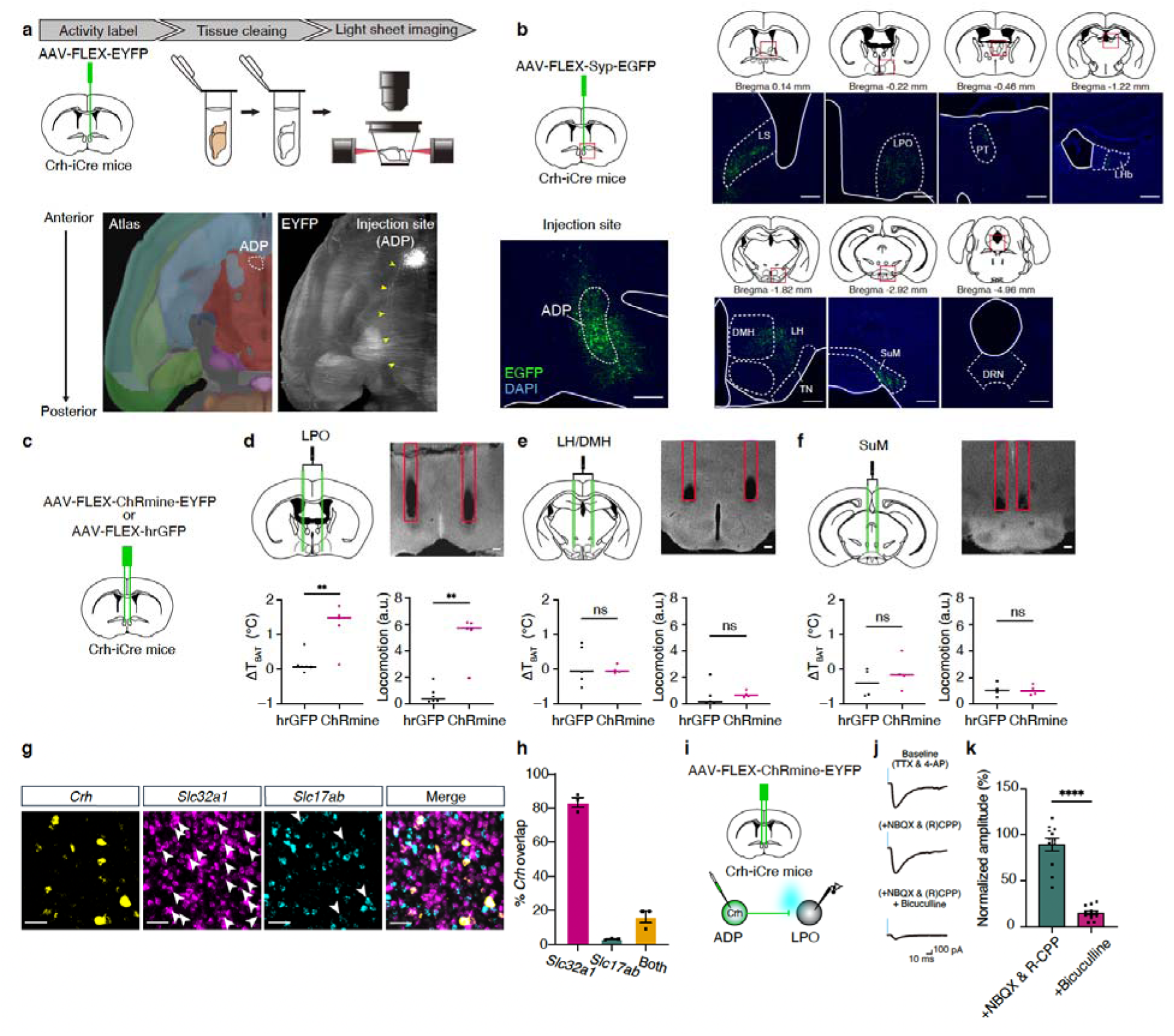
The ADP to LPO pathway promotes thermogenesis and locomotion. **a,** Top, experimental schematic for whole-brain projection mapping of ADP^Crh^ neurons. A Cre-dependent AAV encoding EYFP (AAV-FLEX-EYFP) was injected into the ADP of Crh-iCre mice, followed by brain clearing and light-sheet fluorescence imaging. Bottom, horizontal Allen Brain Atlas section and representative cleared-brain images showing labelled projections. Yellow arrows indicate axons extending posteriorly within the hypothalamus. **b**, Projection mapping of ADP^Crh^ neurons. A Cre-dependent AAV encoding synaptophysin-EGFP (AAV-FLEX-synaptophysin–EGFP) was injected into the ADP of Crh-iCre mice. Representative coronal images show EGFP-labelled axon terminals in downstream regions, including the lateral septum (LS), lateral preoptic area (LPO), lateral habenula (LHb), lateral hypothalamus/dorsomedial hypothalamus (LH/DMH), supramammillary nucleus (SuM), and dorsal raphe nucleus (DRN). The EGFP signal is amplified by anti-GFP immunostaining. Approximate bregma coordinates are indicated above. Scale bars, 300 µm. **c**, Experimental schematic for terminal photostimulation. A Cre-dependent AAV encoding ChRmine (AAV-FLEX-ChRmine-EYFP) or hrGFP control (AAV-FLEX-hrGFP) was injected into the ADP of Crh-iCre mice, and an optical fibre was implanted above the indicated projection target. **d**, Terminal photostimulation in the LPO under euthermic, ad libitum-fed conditions. Top, schematic, and representative image of fibre placement. Bottom, quantification of stimulation-evoked changes in BAT temperature and locomotion (hrGFP, *n* = 6 mice; ChRmine, *n* = 5 mice; two-sided Mann–Whitney *U* test). **e**, Terminal photostimulation in the LH/DMH. Top, schematic and representative image of fibre placement. Bottom, quantification of stimulation-evoked changes in BAT temperature and locomotion (ΔT_BAT_: hrGFP, *n* = 5 mice; ChRmine, *n* = 4 mice; two-sided Welch’s *t*-test; locomotion: hrGFP, *n* = 5 mice; ChRmine, *n* = 4 mice; two-sided Mann–Whitney *U* test). **f**, Terminal photostimulation in the SuM. Top, schematic and representative image of fibre placement. Bottom, quantification of stimulation-evoked changes in BAT temperature and locomotion (hrGFP, *n* = 4 mice; ChRmine, *n* = 4 mice; two-sided unpaired *t*-test). Scale bars, 200 μm. **g**, Representative multiplex in situ hybridization images showing *Crh*, *Slc32a1*, and *Slc17a6* expression in the ADP. Arrowheads indicate *Slc32a1*- or *Slc17a6*-co-expressing *Crh-*positive cells. Scale bar, 50 μm. **h**, Quantification of neurotransmitter phenotype among *Crh*-positive cells, showing overlap with *Slc32a1*, *Slc17a6*, or both. **i**, Experimental schematic for ex vivo electrophysiological analysis of the ADP^Crh^→LPO pathway. A Cre-dependent AAV encoding ChRmine–EYFP (AAV-FLEX-ChRmine-EYFP) was injected into the ADP of Crh-iCre mice, and light-evoked synaptic responses were recorded from LPO neurons in acute brain slices during photostimulation of ADP^Crh^ axon terminals. **j**, Representative traces of light-evoked responses in LPO neurons during pharmacological isolation. Scale bars, 10 ms, 100 pA. **k**, Quantification of normalized light-evoked response amplitude in LPO neurons under the indicated pharmacological conditions (+NBQX & R-CPP versus +Bicuculline; *n* = 12 cells, paired two-sided t-test). Bars indicate mean ± s.e.m.; each dot represents one mouse in **d–f** and one recorded cell in **k**. \*\**P* < 0.01, \*\*\*\**P* < 0.0001; ns, not significant.

To characterize the neurotransmitter phenotype of ADP^Crh^ neurons, we next performed multiplex in situ hybridization. This analysis revealed that ADP^Crh^ neurons are predominantly GABAergic, approximately 90% of *Crh*-positive cells co-expressed *Slc32a1*(Vgat), whereas only ∼15% co-expressed *Slc17a6*(Vglut2) (Fig. 4g, h). To further investigate the synaptic mechanism of the ADP^Crh^ →LPO pathway, we injected a Cre-dependent AAV encoding ChRmine-EYFP into the ADP of Crh-iCre mice. In acute brain slices, photostimulation of EYFP-labelled axon terminals evoked synaptic responses in LPO neurons, which were resistant to tetrodotoxin (TTX) application (12 cells from 7 mice) (Fig. 4i, j). In all LPO neurons receiving light-evoked ADP^Crh^ inputs, these responses persisted in the presence of AMPAR antagonist NBQX and NMDAR antagonist (R)-CPP (Fig. 4j, k). Subsequent application of bicuculline, a GABAA receptor antagonist, dramatically suppressed the responses (Fig. 4j, k). Together, these results demonstrate that ADP^Crh^ neurons exert a direct, monosynaptic inhibitory influence on downstream LPO neurons via GABA release.

Finally, to assess whether the recruitment of ADP^Crh^ neurons during rewarming is conserved in a hibernating species, we examined the Syrian hamster (*Mesocricetus auratus*). First, we confirmed that *Crh*-expressing neurons are present in the ADP of Syrian hamsters (Extended Data Fig. 9a). We next collected brains from hamsters that had spontaneously recovered to a body temperature of 25-30 °C after days of deep torpor (Extended Data Fig. 9b). Multiplex in situ hybridization revealed broad *c-Fos* induction in the hypothalamus, and approximately 64.51 % of *Crh*-expressing cells in the ADP co-expressed *c-Fos*. In contrast, euthermic control brains showed little *c-Fos* expression in this region (Extended Data Fig. 9c, d). Thus, recruitment of ADP^Crh^ neurons during rewarming is not unique to mice but is also observed in a hibernating species.

## Discussion

In this study, we identify ADP^Crh^ neurons as a key population promoting rewarming from fasting-induced torpor in mice. Loss- and gain-of-function manipulations show that these neurons limit torpor depth and promote timely recovery to euthermia by engaging BAT thermogenesis, which precedes locomotor arousal. This pathway acts through a GABAergic projection to the lateral preoptic region and is recruited during rewarming in a hibernating species, suggesting conserved logic for exiting deep torpor.

ADP^Crh^ neurons were dispensable for acute cold defence, stress hyperthermia and LPS-induced fever, yet were required to limit torpor depth and duration. Canonical models place thermogenic control within a POA–DMH–RPa axis^17^, and several studies have implicated the DMH in torpor induction^9,18,19^. Yet projection-specific stimulation in our study identified the LPO, but not the DMH or LH, as a sufficient downstream target of ADP^Crh^ neurons for promoting BAT-mediated thermogenesis. Together, these findings suggest that torpor rewarming is not simply mediated by re-engagement of a general thermoregulatory pathway. Instead, the ADP^Crh^→LPO projection appears to selectively support timely exit from torpor. Whether this pathway ultimately converges on classical DMH- and/or RPa-dependent thermogenic effectors remains unresolved.

The LPO contains intermingled populations implicated in torpor and sleep-related cooling, including avMLPA^Adcyap1^neurons^5^ and subsets of galanin neurons^20,21^. Although the postsynaptic targets remain unknown, ADP^Crh^ inputs may promote rewarming by inhibiting LPO pathways that stabilize hypothermia and quiescence, thereby permitting rapid thermogenesis and behavioural arousal.

Rewarming is initiated despite a global reduction in neural activity during torpor. Therefore, the circuitry that drives rewarming must remain excitable at low body temperature, and an appropriate trigger must be able to engage it. One possibility is that ADP^Crh^ neurons are activated by ascending subcortical pathways that convey thermosensory information to the preoptic area^22^. Brainstem catecholaminergic neurons innervate the preoptic region and have been implicated in torpor induction^7^, suggesting that these inputs may modulate the activity of ADP^Crh^ neurons during rewarming. Identifying the presynaptic inputs to ADP^Crh^ neurons via retrograde trans-synaptic tracing will be important for testing this idea. In addition to neural inputs, circulating energy-balance signals may modulate the excitability of ADP^Crh^ neurons. Leptin suppresses torpor^23,24^, whereas ghrelin can deepen torpor bouts^25^, and the relatively permeable blood–brain barrier of the hypothalamus raises the possibility of direct actions on ADP neurons. However, as these hormones are unlikely to exhibit abrupt concentration changes during a single bout, they may act as permissive gates rather than acute triggers of rewarming.

Together, our findings identify a preoptic circuit that promotes recovery from an extreme energy-saving state and suggest similar circuit logic in a hibernating species.

---

**Extended Data Fig. 1:**
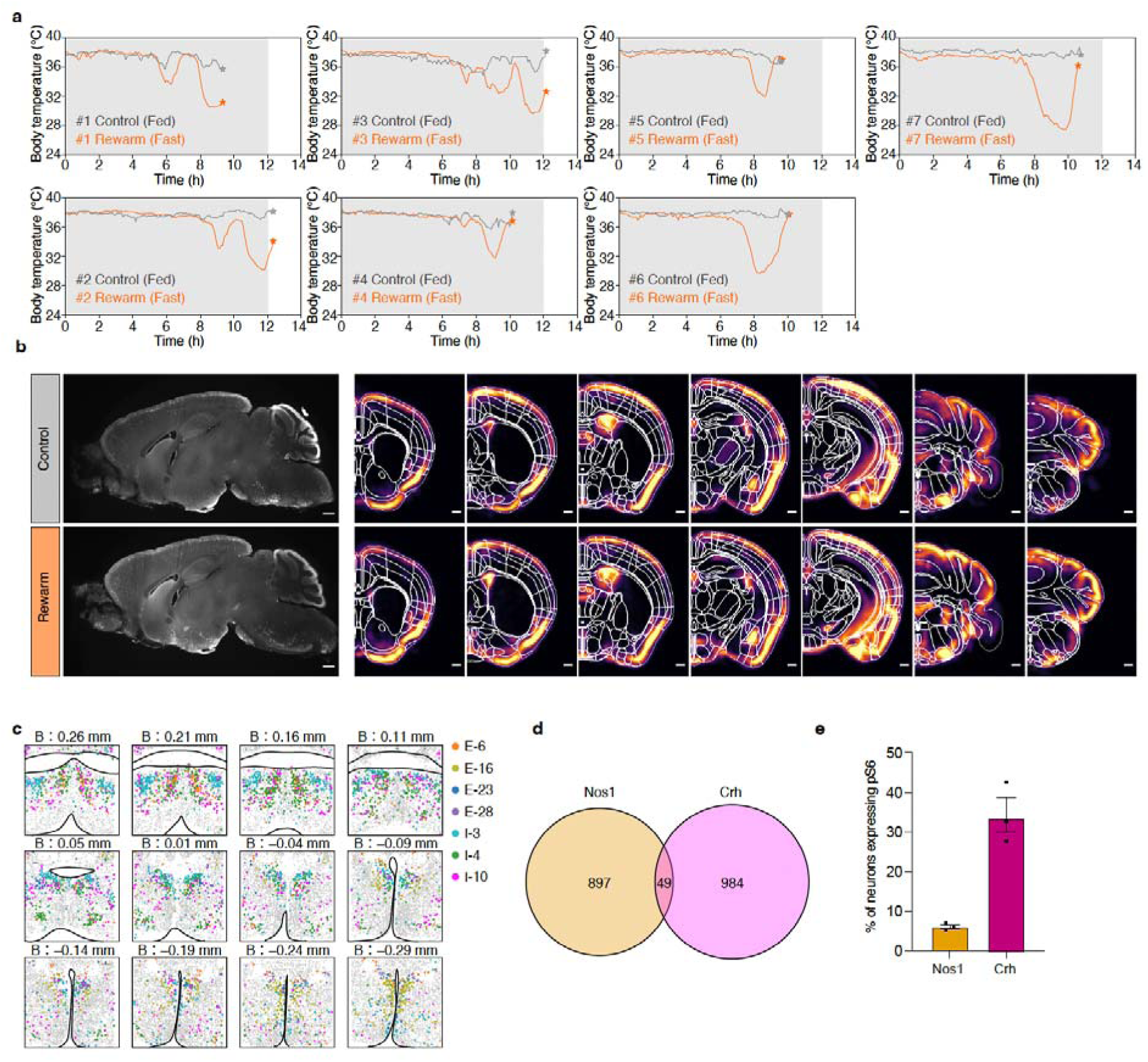
Additional data for the identification of rewarming-associated regions. **a**, Body temperature traces of individual mice used for whole-brain screening. Grey shading indicates the dark phase; asterisks indicate the perfusion time. **b**, Representative light-sheet image of cleared brain sagittal section showing pS6 signal (left) and heatmaps of mean pS6 signal density along the rostrocaudal axis (right). Scale bars, 500 µm. **c,** Spatial distribution of ADP-enriched clusters in the published atlas, referred to as the anterior parvicellular paraventricular hypothalamic nucleus (PaAP) in Moffitt et al^13^. **d**, Venn diagram showing the overlap between *Nos1*- and *Crh*-positive neurons within the ADP (*n* = 3 mice). **e**, Fraction of pS6-positive neurons in *Nos1*- or *Crh*-positive neurons in rewarming brains.

**Extended Data Fig. 2:**
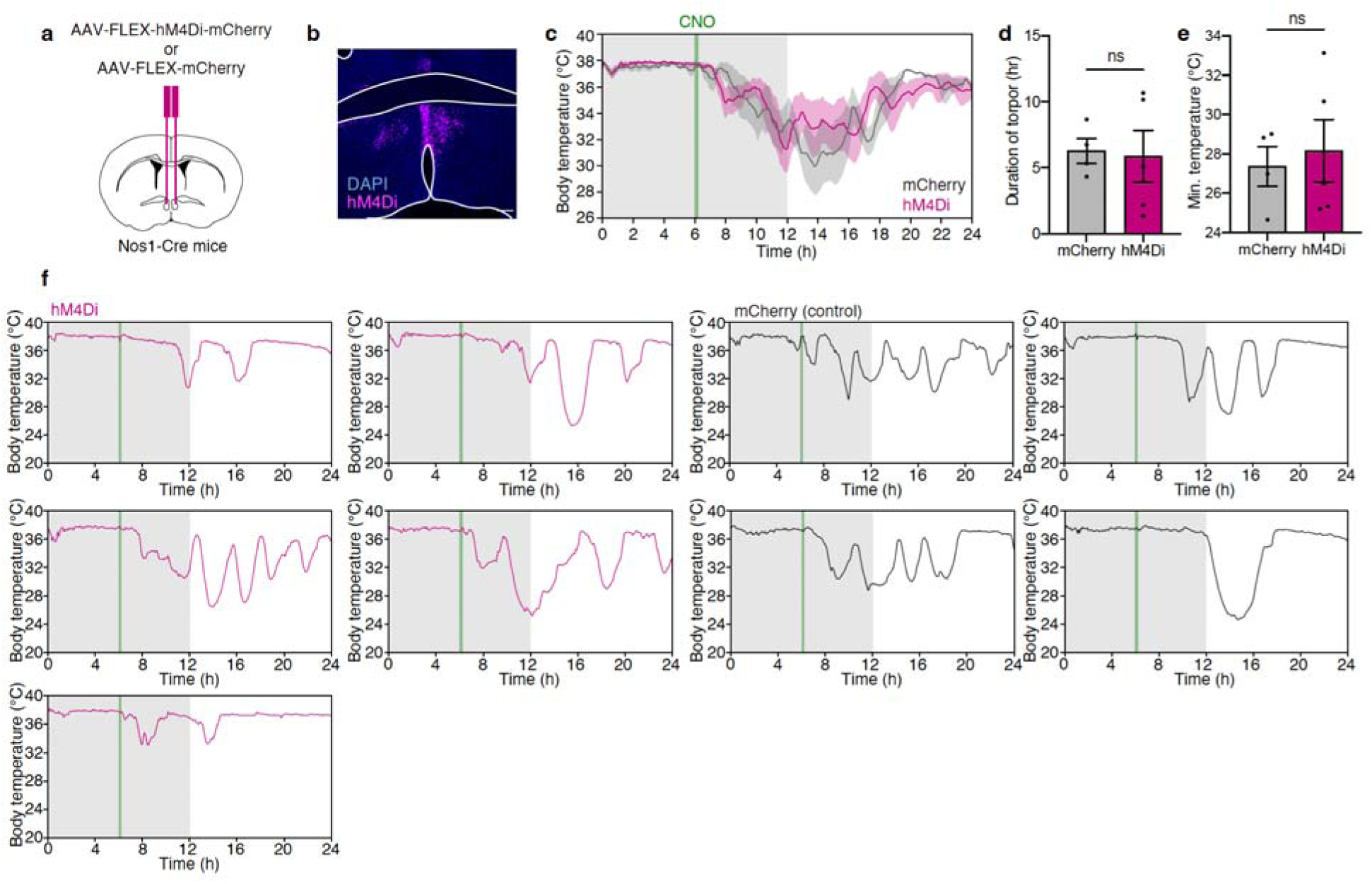
MnPO/ADP-Nos1 neurons are not required for rewarming from torpor. **a**, Experimental schematic for chemogenetic inhibition. A Cre-dependent AAV encoding hM4Di–mCherry (AAV-FLEX-hM4Di-mCherry) or mCherry control (AAV-FLEX-mCherry) was bilaterally injected into the ADP region of Nos1-Cre mice. **b**, Representative fluorescence image showing viral expression in the MnPO/ADP region. Scale bar, 200 µm. **c**, Body temperature during 24-hour fasting with CNO administration. Fasting was initiated at ZT12, and CNO was injected 6-hour after fasting onset (green line). Grey shading indicates the dark phase. Traces show mean ± s.e.m.. **d**, Torpor duration (mCherry, *n* = 4 mice; hM4Di, *n* = 5 mice; two-sided unpaired *t*-test) **e**, Minimum Tb (mCherry, *n* = 4 mice; hM4Di, *n* = 5 mice; two-sided unpaired *t*-test). **f,** Body temperature traces from individual mice in the hM4Di (magenta) and mCherry control (black) groups. Bars indicate mean ± s.e.m. in **d,e**. ns, not significant.

**Extended Data Fig. 3:**
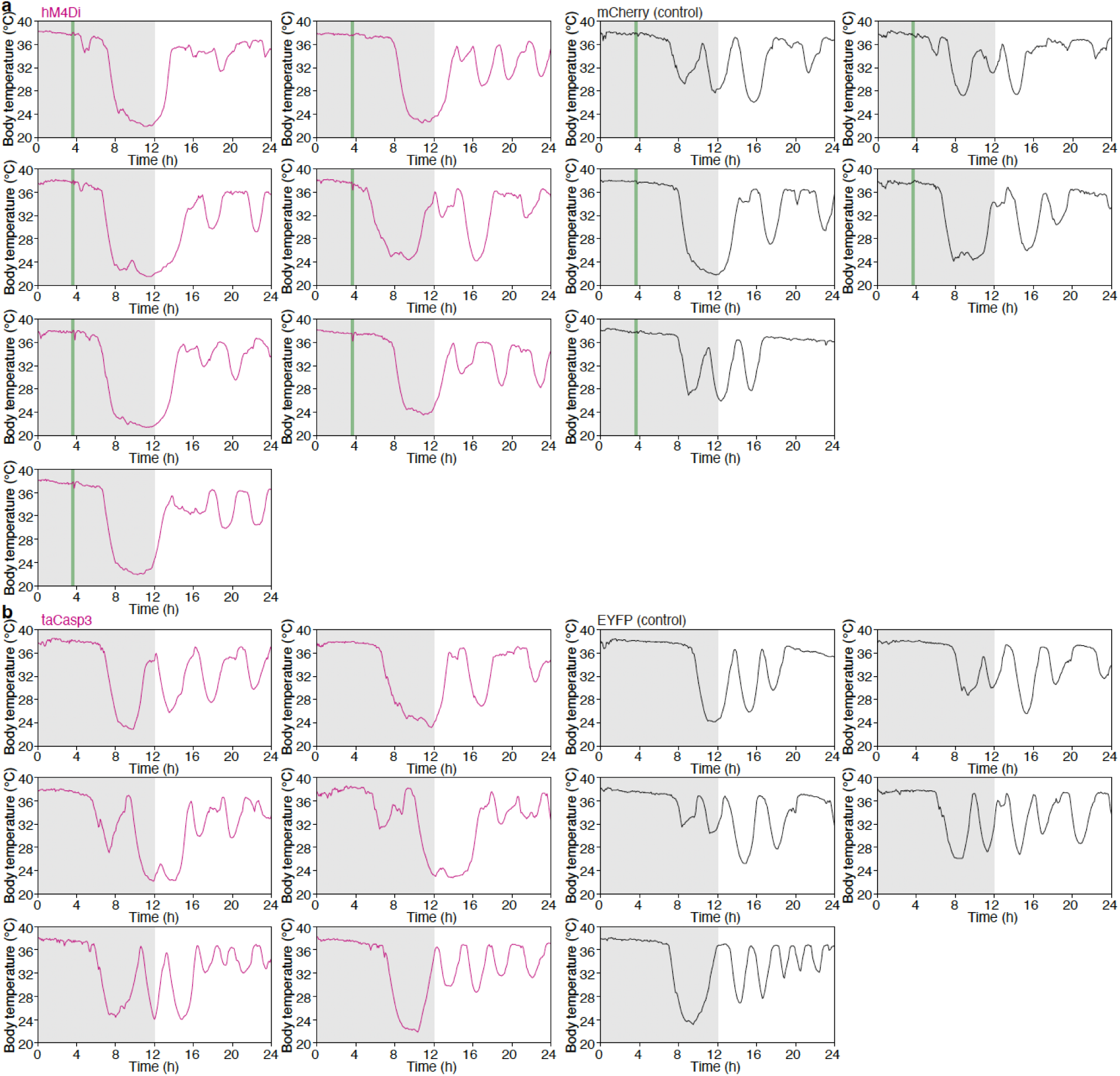
Additional data on loss-of-function experiments of ADP^Crh^ neurons. **a**, Body temperature traces from individual mice in the hM4Di (magenta) and the mCherry control (black) groups during 24-h fasting. Grey shading indicates the dark phase; green line indicates the timing of CNO injection. **b**, Body temperature traces from individual mice in the taCasp3 (magenta) and EYFP control (black) groups during 24-h fasting. Grey shading indicates the dark phase.

**Extended Data Fig. 4:**
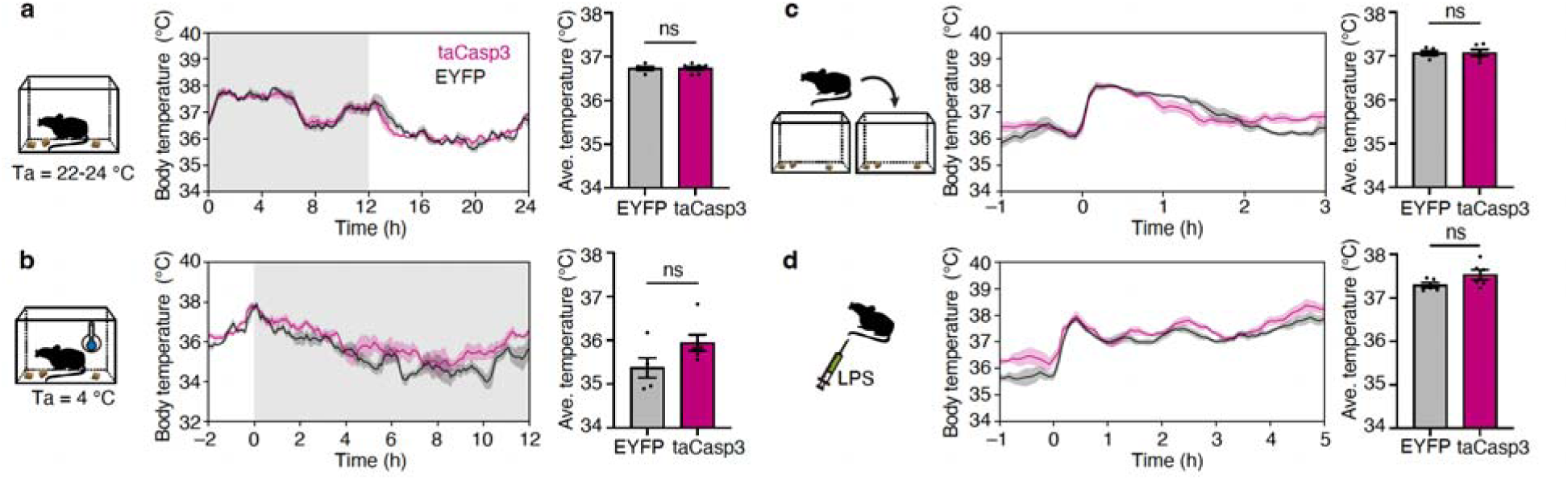
Effects of ablation on LPS, stress, cold-induced thermogenesis. **a**, Body temperature traces under ad libitum-fed conditions at ambient temperature (Ta = 22–24 °C) in EYFP control (n = 5 mice) and taCasp3 (n = 6 mice) (left), with quantification of mean body temperature (right; two-sided unpaired *t*-test). Traces show the mean across 4 days. Grey shading indicates the dark phase. **b**, Body temperature traces during cold exposure (Ta = 4 °C; left), with quantification of mean body temperature for 12 h cold exposure (right; two-sided Mann–Whitney *U* test). Grey shading indicates the dark phase. **c**, Body temperature changes in response to cage-change stress (left), with quantification of mean body temperature during the 3h after cage changes (right; two-sided unpaired t-test). **d**, Body temperature response to LPS administration (50 µg kg ¹) (left), with quantification of mean body temperature during the 5 h after injection (right; two-sided unpaired t-test). Bars indicate mean ± s.e.m.; ns, not significant.

**Extended Data Fig. 5:**
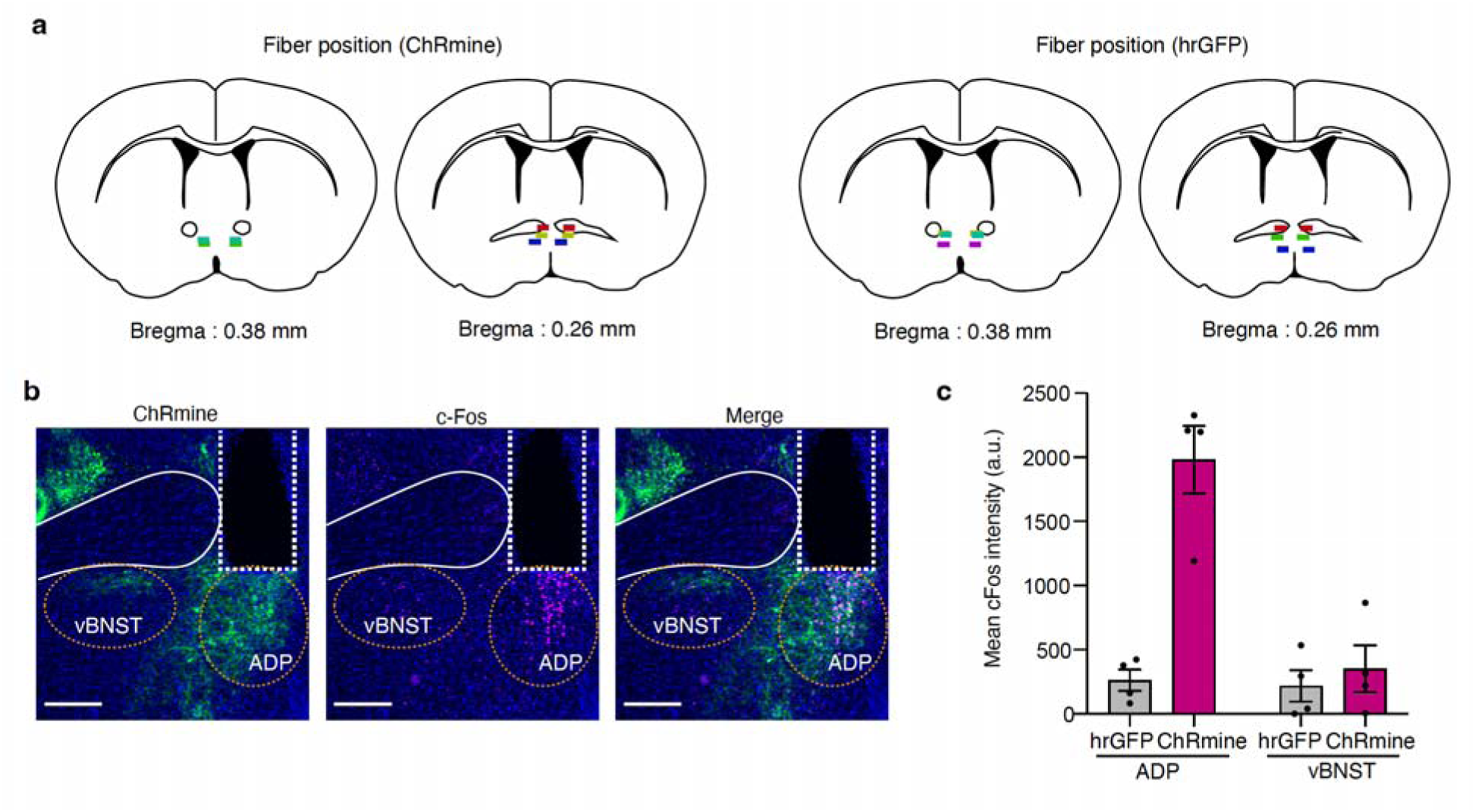
Validation of regional specificity of optogenetic activation. **a**, Schematic showing optic fibre placements for ChRmine and hrGFP control groups. Each colour denotes an individual mouse. **b**, Representative fluorescence images showing c-Fo immunofluorescence induced by photostimulation in the ADP, with no evident activation in the ventral BNST (vBNST). Circled regions indicate the regions of interest used for quantification. Scale bar, 100 µm. **c**, Quantification of mean c-Fos intensity in the ADP and vBNST following photostimulation in ChRmine and hrGFP mice. Bars indicate mean ± s.e.m.

**Extended Data Fig. 6:**
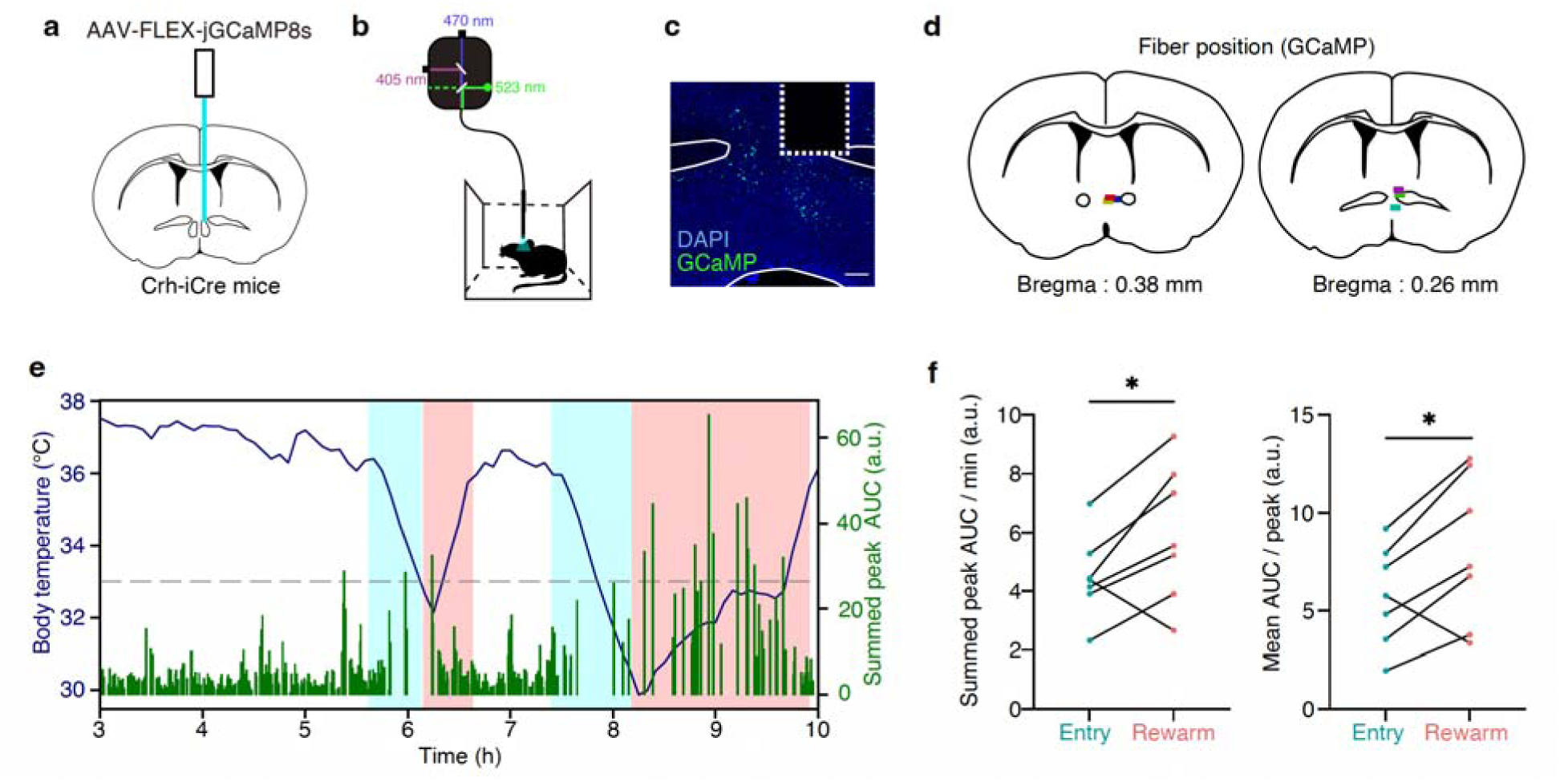
Neural dynamics of ADP^Crh^ neurons during torpor. **a**, Experimental schematic for fibre photometry. A Cre-dependent AAV encoding jGCaMP8s (AAV-FLEX-jGCaMP8s) was injected into the ADP of Crh-iCre mice, and an optical fibre was implanted above the ADP. **b,** Schematic of the fibre photometry recording configuration. **c**, Representative fluorescence image showing jGCaMP8s expression in the ADP and fibre trace. Scale bar, 200 µm. **d**, Fibre placements for photometry recordings. Each colour denotes an individual mouse. **e**, Representative recording showing body temperature (navy) and summed peak calcium-event AUC (green) during fasting-induced torpor. Light blue and red shading indicate torpor-entry and rewarming phases, respectively. **f**, Quantification of average peak AUC during torpor entry and rewarming (*n* = 12 bouts from 7 mice, paired two-sided t-test). **g,** Quantification of summed peak AUC per minutes (left) and mean AUC per peak (right) during torpor entry and rewarming (*n* = 7 mice, paired two-sided t-test). \**P* < 0.05.

**Extended Data Fig. 7:**
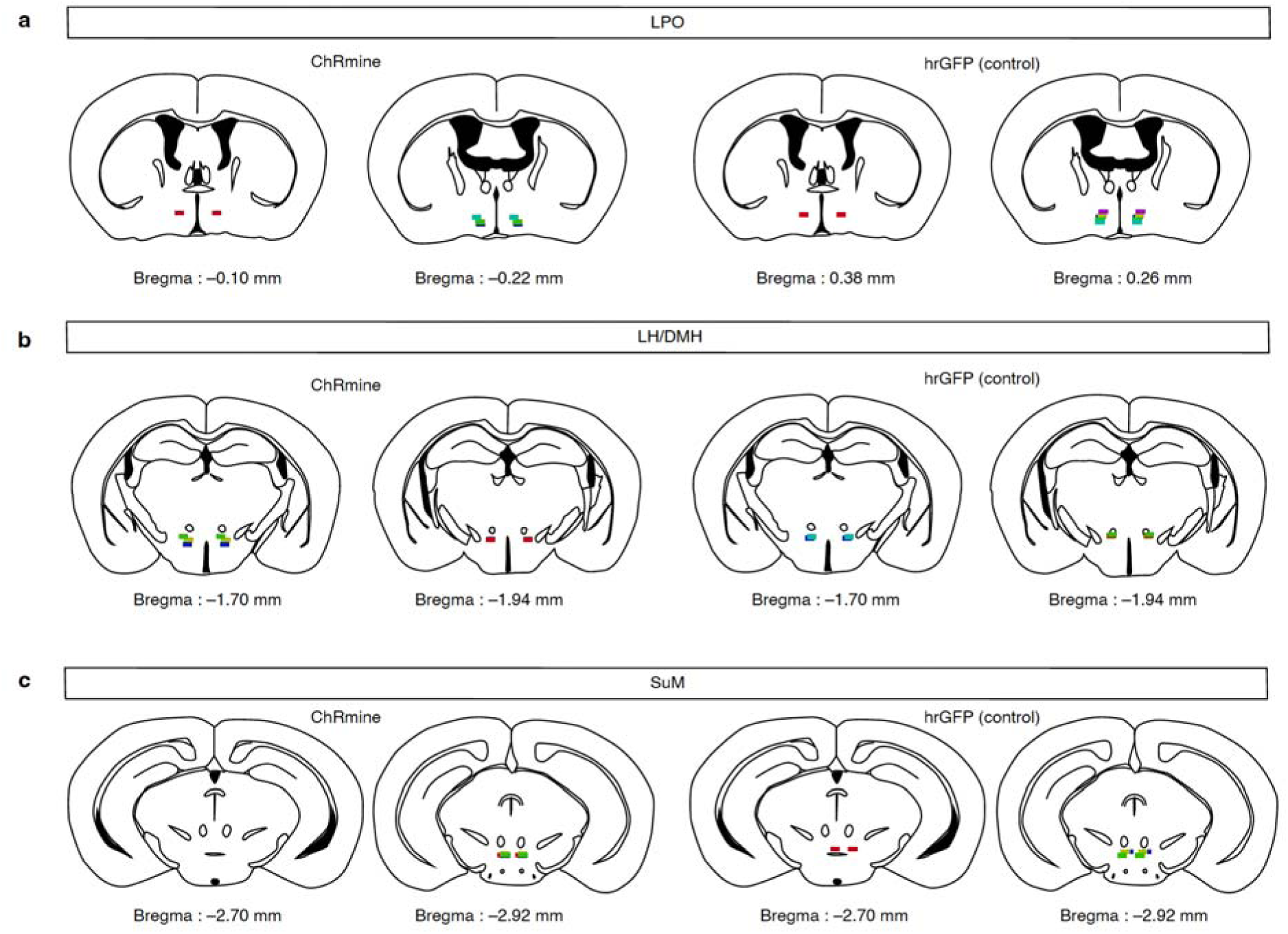
Validation of tip location for terminal stimulation. Schematic showing optic fibre tip locations in ChRmine and hrGFP control mice for terminal stimulation in the LPO (**a**), LH/DMH (**b**) and SuM (**c**). Each colour denotes an individual mouse.

**Extended Data Fig. 8:**
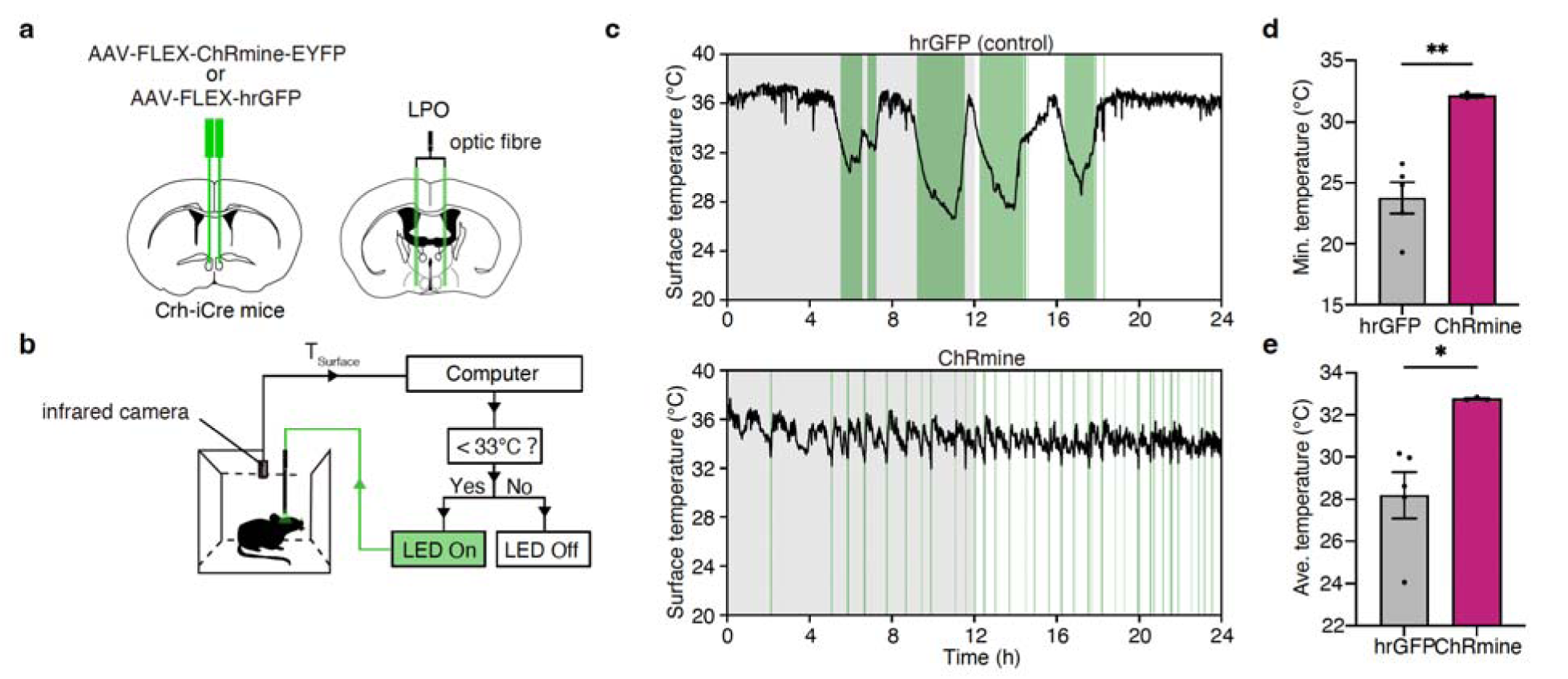
Closed-loop activation of ADP^Crh^→LPO pathway trigger rewarming. **a,** Experimental schematic for terminal photostimulation. A Cre-dependent AAV encoding ChRmine (AAV-FLEX-ChRmine-EYFP) or hrGFP control (AAV-FLEX-hrGFP) was injected into the ADP of Crh-iCre mice, and an optical fibre was implanted above the LPO. **b**, Closed-loop optogenetic stimulation paradigm during fasting. Surface temperature was monitored in real time by an infrared camera, and stimulation was triggered when the temperature fell below 33 °C. **c**, Representative surface temperature traces during fasting with closed-loop terminal stimulation of ADP^Crh^→LPO axons in hrGFP control (top) and ChRmine mice (bottom). Green shading indicates epochs of light delivery. Grey shading indicates the dark phase. **d**, Minimum surface temperature during the 24-hour fasting (hrGFP, *n* = 5 mice, ChRmine, *n* = 3 mice; two-sided unpaired *t*-test with Welch’s correction). **e**, Mean surface temperature during photostimulation period (hrGFP, *n* = 5 mice; ChRmine, *n* = 3 mice; two-sided unpaired *t*-test with Welch’s correction). Bars indicate mean ± s.e.m.; \**P* < 0.05, \*\**P* < 0.01.

**Extended Data Fig. 9:**
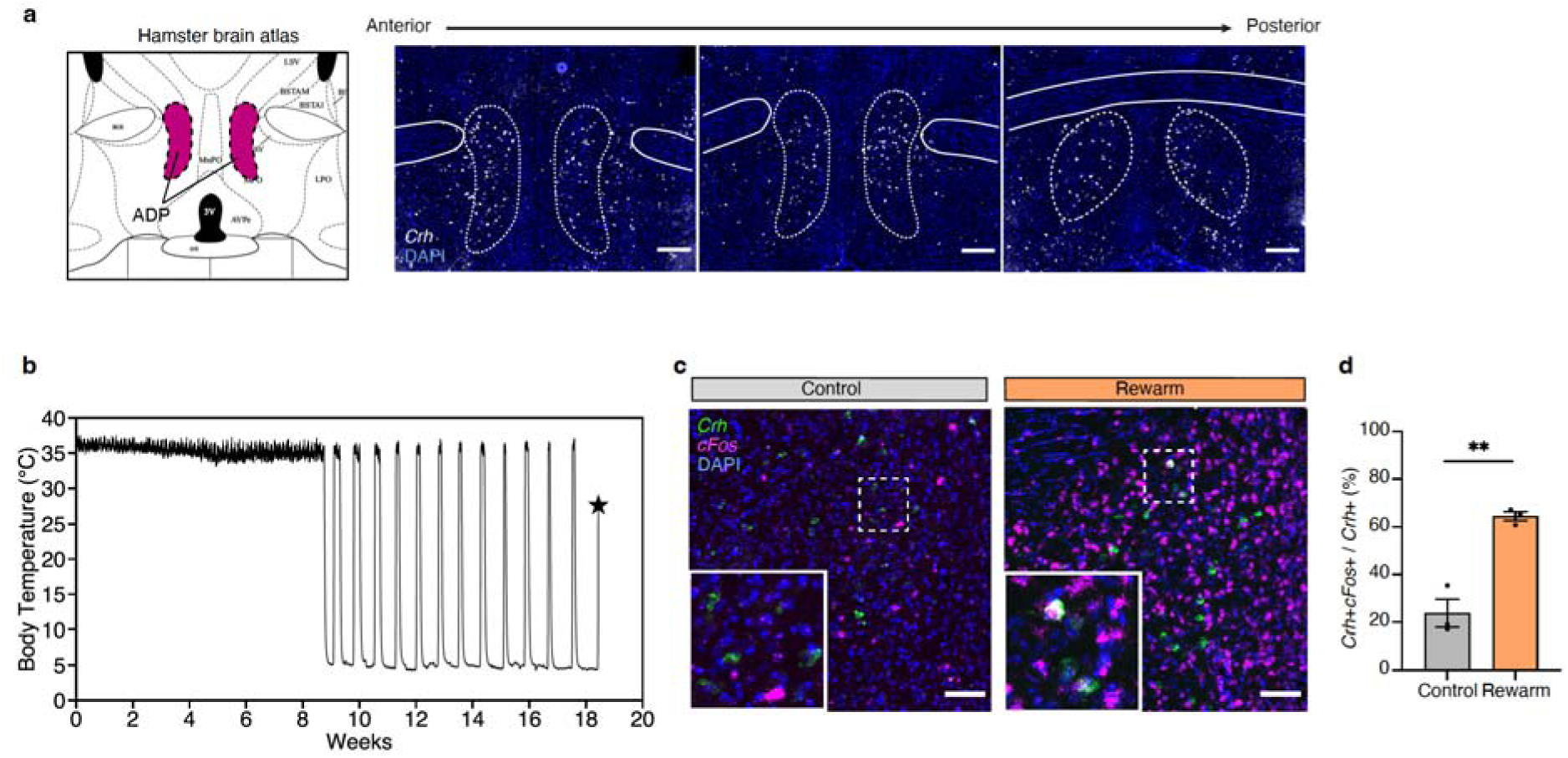
ADP^Crh^ neurons are recruited during rewarming in a hibernating species. **a,** The hamster brain atlas at bregma + 0.9 mm (left) and representative fluorescence images showing *Crh*-positive cells around the ADP. Scale bar, 300 µm. **b,** Representative body temperature trace of a hibernating hamster. Week 0 indicates the onset of exposure to winter-like conditions (4–5 °C with a 8 h/16 h light-dark cycle) for hibernation induction. The control group was continuously housed under summer-like conditions (24–25 °C with a 14/10 h light-dark cycle). Brains were collected from Syrian hamsters that had spontaneously rewarmed to a body temperature of 25–30 °C after several days of deep hibernation. The star indicates the time of perfusion. **c,** Representative fluorescence images showing *c-Fos*-positive *Crh* neurons in the ADP region under control (left) and rewarm (right) conditions. Scale bar, 100 µm. **d,** Quantification of the percentage of *c-Fos*-positive cells among *Crh*-positive cells in the ADP of control and rewarming hamsters. The mean percentage was 23.75% in controls and 64.51% in rewarming hamsters (Control, *n* = 3; Rewarm, *n* = 3). Bars indicate mean ± s.e.m. Two-tailed unpaired t-test. \*\**P* < 0.01.

## Methods

### Mice

Wild-type C57BL/6J mice were obtained from CLEA Japan. Crh^tm2(icre)ksak^ (Crh-iCre) were kindly provided by Dr. Kenji Sakimura. B6.Cg-Gt(ROSA)26Sortm14(CAG-tdTomato)Hze/J(Ai14) and B6.129-Nos^1tm1(cre)Mgmj^/J(Nos1-Cre) mice were obtained from Jackson Laboratories. Unless otherwise stated, all Cre-driver mice used in this study were heterozygous females aged 12–20 weeks. Mice were housed in Plexiglas chambers under a reversed light-dark cycle (lights on from 08:00 to 20:00 or from 06:00 to 18:00) at 22 ± 1 °C (unless otherwise indicated), with food and water available ad libitum. All experimental procedures were approved by the Animal Care and Use Committee of Nagoya University (Approval No.: R240004) and the National Institute for Physiological Sciences (Approval No.: 25A065, 26A060). All efforts were made to reduce the number of animals used and to minimize animal suffering.

### Torpor induction

To record core body temperature, mice were anaesthetized with isoflurane and intraperitoneally implanted with a temperature logger (Star-Oddi, DST nano-T). After a recovery period of at least 14 days following surgery, mice were transferred from their home cage to a new cage in a temperature-controlled room (16) at the onset of the dark period. The mice were then fasted, or ad libitum fed for 24 hours. Water was available ad libitum for both groups throughout the experiment. Data on core body temperature were collected and analysed by Mercury software (Star-Oddi).

### Brain sampling from mice during torpor rewarming

Wild-type C57BL/6J female mice were fasted or ad libitum fed at 16 as described above. To determine the nadir (minimum body temperature) in real time, body surface temperature was continuously monitored at 1-min intervals using an infrared thermal imaging camera (Optris PI450). For rewarming screening, bouts in which body surface temperature remained below 28 °C for at least 5 min were selected for nadir detection. Mice were deeply anaesthetized with isoflurane and transcardially perfused with PBS followed by 4% paraformaldehyde (PFA), and brains were collected 30 to 60 minutes after reaching the nadir, corresponding to the period of active rewarming. To strictly control for circadian variables, a fed control mouse was temporally paired with each fasted mouse and processed at the exact matched time point.

### Brain sampling from Syrian Hamsters during torpor rewarming

4- or 8-week-old male Syrian hamsters were purchased from an outbred colony (Japan SLC., Inc) and housed under summer-like long-photoperiod and warm conditions (24–25 °C with a 14/10 h light-dark cycle) with ad libitum access to water and food (MR standard diet, Nihon Nosan). 3–4 individuals were housed per cage, and their body mass was recorded weekly when the cages were replaced.

At 12 weeks old, all animals were transferred to individual polypropylene cages with ad libitum access to water and food. The control group was continuously housed under summer-like conditions after that. The TB logger (iButton, #DS1925L-F5, Maxim Integrated), calibrated by the manufacturer, was coated with rubber (total mass approx. 3.5 g; Plasti Dip, Performix Coatings) and surgically implanted into the abdominal cavity of hamsters to induce hibernation under 2% isoflurane anaesthesia. The logger recorded Tb every 10 min with a resolution of 0.0625 °C. After surgery, animals were allowed to recover for 2 weeks.

For hibernation induction, the animals were transferred to winter-like short-photoperiod and cold conditions (4–5 °C with an 8/16 h light-dark cycle) with ad libitum access to water and food. Body mass was measured weekly, and cages were replaced every other week, except for hibernating animals. 2–3 months after the transfer to winter-like conditions, animals began to hibernate and displayed typical torpor-arousal cycles.

For collecting brain samples during the rewarming phase from deep torpor, hibernating hamsters were set to cages in which the oxygen consumption rate of the animals was monitored in real-time using the respiratory gas analyser ARCO-2000 (ARCO System). 2 hours after observing a significant increase in oxygen consumption, we monitored the oral temperature of rewarming animals every 10 min using a wired temperature meter (BWT-100A, Bioresearch center). When the detected oral temperature reached 30 °C (while, the iButton recorded Tb of 25–30 °C), the animals were immediately perfused with PBS, followed by 4% PFA/PBS, under 2% isoflurane anaesthesia. After perfusion, Tb records were retrieved from the iButton, and the brains were collected and further fixed in 4 %PFA/PBS for 1 day at 4 °C. The fixed brains were then dehydrated in 30% sucrose/PBS for 2 days at 4 °C. Finally, the dehydrated brains were embedded in O.C.T. compound (Sakura Tissue-Tek), flash-frozen, and stored in –80 °C until sectioning. Brains of the control group were collected in parallel and processed in the same way as those from the rewarming animals.

### Whole-brain immunolabeling and iDISCO clearing

Whole-brain clearing and immunolabelling were performed following the iDISCO protocol, as previously described^9^. Fixed mouse brains were dehydrated in a graded methanol series, delipidated in dichloromethane (DCM)(nacalai tesque, 22414-23)/methanol, and bleached with 5% H_2_O_2_ in methanol overnight at 4 °C. After rehydration, samples were permeabilized and blocked, followed by incubation with the primary antibody (rabbit anti-phospho-S6, 1:300; Invitrogen, 44923G) for 7 days at 37 °C. After extensive washing, samples were incubated with the secondary antibody (Alexa Fluor 647 donkey anti-rabbit IgG, 1:500; Invitrogen, A31573) for 7 days at 37 °C. Finally, immunolabelled brains were dehydrated, washed in DCM, and cleared in dibenzyl ether (Merck, 108014).

### Light-sheet fluorescence imaging for whole-brain activity mapping

Optically cleared brain samples were imaged sagittally (right lateral side up) on a light-sheet fluorescence microscope (Miltenyi Biotec, Ultramicroscope II) equipped with a CMOS camera and a ×0.63 objective lens. Volume scanning was performed at a 5-μm step size using a continuous light-sheet scanning method with the included contrast-mixing algorithm for the 640 nm and 488 nm channels.

### Automated regional quantification of whole-brain activity

Phospho-S6-immunoreactive cells were automatically detected and registered using the ClearMap software pipeline, as previously described^9^. Briefly, background subtraction was performed using a disk-shaped structural element (7-pixel diameter) and morphological opening, followed by 3D peak detection with a cell-size threshold of 700 voxels. For spatial normalization, autofluorescence volumes were registered to the Allen Mouse Brain Reference Atlas (25 μm/voxel) utilizing the average STPR brain template^26^. To generate whole-brain pS6 heatmaps, each detected pS6-positive cell was computationally represented as a uniform sphere with a diameter of 375 μm. Regional pS6 signal intensity was then quantified from these heatmaps on the basis of atlas-defined brain regions and compared between groups. Group differences in regional pS6 signal intensity were assessed using two-sided Welch’s t-tests followed by Benjamini–Hochberg false discovery rate correction, and fold changes were calculated for each region. Voxel-wise t-tests were additionally performed to generate p-value maps for visualization.

### Viral constructs

The following AAV vectors were used in this study: AAV9-CAG-FLEX-hM4Di-mCherry (4.7x10^12^ vg mL^−1^; in-house preparation), AAV9-CAG-FLEX-mCherry (1.8x10^12^ vg mL^−1^; in-house preparation), AAV9-CAG-FLEX-ChRmine-EYFP (7.2x10^12^vg mL ^− 1^; in-house preparation), AAV9-CMV-FLEX-hrGFP (4.0x10^12^vg mL^−1^; in-house preparation), AAV2-EF1α-DIO-EYFP (4.5x1012 vg mL^−1^; UNC Vector Core), AAV2-CAG-FLEX-synaptophysin–EGFP (4.8x10^12^ vg mL^− 1^; Brain/MINDS 2.0 vector core), AAV9-CAG-FLEX-jGCaMP8s (4.7x10^13^ vg mL 1; in-house preparation), and AAV2-FLEX-taCasp3-TEVp (4.3x10^12^ vg mL^−1^; UNC Vector Core).

### MERFISH data analysis

A published spatial transcriptomic atlas was obtained from Dryad (doi:10.5061/dryad.8t8s248), corresponding to the dataset reported by Moffitt et al^13^. To identify clusters enriched in the ADP, neuronal clusters annotated to the anterior parvicellular paraventricular hypothalamic nucleus (PaAP), which corresponds to the ADP in this atlas, were selected based on the published annotation. This yielded three inhibitory clusters (I-3, I-4 and I-10) and four excitatory clusters (E-6, E-16, E-23 and E-28). Among these seven ADP-enriched clusters, three clusters (E-6, I-4, and I-10) were selected for further analysis based on their spatial similarity to the distribution of rewarming-associated pS6-positive cells. For each target cluster, gene enrichment was assessed by comparing “gene expression values” in neurons belonging to that cluster with that in all other neurons. For each gene, cluster enrichment was quantified as the log_2_ fold change based on mean expression. Statistical significance was assessed using a two-sided Mann–Whitney *U* test, and *P* values were corrected for multiple comparisons using the Benjamini–Hochberg false discovery rate procedure.

### Immunohistochemistry

Mice were anaesthetized with isoflurane and transcardially perfused with PBS followed by 4% PFA. Brains were collected, post-fixed in 4% PFA at 4 °C overnight, and cryoprotected in 30% sucrose in PBS at 4 °C for at least 2 days. Coronal brain sections (60 μm or 40 μm) were prepared using a Leica CM3050S cryostat (Leica Microsystems) or RWD Minux FS800A cryostat. Free-floating sections were blocked for 1 h in blocking buffer consisting of 3% bovine serum albumin in PBS containing 0.3% Triton X-100, and then incubated with primary antibodies overnight at room temperature. After washing in PBS containing 0.3% Triton X-100, sections were incubated with secondary antibodies for 5 h at room temperature. Primary antibodies used in this study were chicken anti-GFP (1:1000, Aves Labs, GFP-1010), rabbit anti-phospho-S6 (1:1,000, Invitrogen, 44-923G), mouse anti-NOS1 (1:100, Santa Cruz Biotechnology, sc-5302), rabbit anti-CRF (1:1,000, BMA Biomedicals, T-4037), and rabbit anti-c-Fos (1:1000, Abcam, ab222699). For double immunofluorescence, rabbit anti-phospho-S6 or rabbit anti-CRF was combined with mouse anti-NOS1 in separate experiments. Secondary antibodies were diluted 1:1,000 in blocking buffer and included CF 488-conjugated donkey anti-chicken IgY (Biotium, 20166), CF 488-conjugated donkey anti-rabbit IgG (Biotium, 20015), CF 594-conjugated donkey anti-rabbit IgG (Biotium, 20152), CF 647-conjugated donkey anti-rabbit IgG (Biotium, 20047), and Alexa Fluor Plus 647-conjugated donkey anti-mouse IgG (Invitrogen, A32787).

### Fluorescence in situ hybridization

For mouse experiments, brains were collected and processed as described in the immunohistochemistry section. Brains were frozen in embedding medium at −80 °C, sectioned at 20 µm using a cryostat, and mounted on glass slides. Multiplex fluorescence in situ hybridization was performed using the RNAscope Multiplex Fluorescent v2 Assay (Advanced Cell Diagnostics, 323100) according to the manufacturer’s instructions. Briefly, sections were pretreated with hydrogen peroxide for 10 min at room temperature, followed by target retrieval at 98–102 °C for 5 min, and Protease III digestion at 40 °C for 30 min in a HybEZ oven (Advanced Cell Diagnostics). Sections were then hybridized with target probe Sections were then hybridized for 2 h at 40 °C with target probes against *Crh* (Advanced Cell Diagnostics, 316091-C1), *Slc32a1* (Advanced Cell Diagnostics, 319191-C4), and *Slc17a6* (Advanced Cell Diagnostics, 319171-C3). Following sequential signal amplification steps, transcript signals were developed using the TSA Plus Fluorescence system with Opal 520 (FP1487A), Opal 570 (FP1488A), and Opal 690 (FP1497A) fluorophores (PerkinElmer, Waltham, MA). Slides were mounted using ProLong Gold Antifade Mountant (Thermo Fisher Scientific, P36930). For spatial mapping and quantification, preoptic area sections were analysed at 160-µm intervals (every eighth section) from each animal (*n* = 3 mice). Images were obtained using a BZ-9000 fluorescence microscope (Keyence).

For hamster experiments, the same procedure was used except that brains were sectioned at 25 µm and hybridized with probes against Crh (Advanced Cell Diagnostics, 1139271-C1) and Fos (Advanced Cell Diagnostics, 516241-C2). Transcript signals were developed using Opal 520 (FP1487A) and Opal 570 (FP1488A). For spatial mapping and quantification, sections were stained and imaged at 100-µm intervals (every fourth section) from each animal (*n* = 3 hamsters). Images were obtained using a BZ-9000 fluorescence microscope (Keyence).

### Cell quantification

To quantify labelled cells, coronal brain sections containing the ADP region were prepared and processed as described above. Z-stack fluorescence images were acquired using an LSM980 laser-scanning confocal microscope (Carl Zeiss) or a BZ-9000 microscope (Keyence) equipped with a 10× or 20× objective lens, and maximum-intensity projections were generated for analysis. Image processing and cell counting were performed using ImageJ/Fiji software (NIH). The anatomical boundaries of the ADP were delineated utilizing the Allen Mouse Brain Reference Atlas as a spatial reference. Immunofluorescence-positive cells within the targeted regions were manually quantified.

### Chemogenetic silencing of ADP neurons

Female heterozygous Crh-iCre or Nos1-Cre mice received bilateral stereotaxic injections of AAV9-CAG-FLEX-hM4Di-mCherry into the ADP (anterior-posterior (AP), +0.35 mm; medial-lateral (ML), ±0.3 mm; dorsal-ventral (DV), –4.9 mm). Control mice received identical AAV9-CAG-FLEX-mCherry injections. Two weeks after the injection, temperature data loggers (Star-Oddi, DST nano-T) were implanted intraperitoneally, and the mice were allowed to recover for an additional two weeks. To induce torpor, mice were housed at 16 °C, and food was removed at the onset of the dark phase. Clozapine-N-oxide (CNO; Enzo Life Sciences, BML-NS105) was administered intraperitoneally at a dose of 1.0 mg kg^-1^ body weight at 3.5 h after fasting onset in Crh-iCre experiments and at 6 h after fasting onset in Nos1-Cre experiments. CNO was initially dissolved in sterile water to yield a 10 mg ml^-1^ stock solution, then diluted in saline to a final working concentration of 100 µg ml^-1^ before injection. One mouse showing unilateral expression due to a missed injection was excluded from the analysis.

### in vivo optogenetic stimulation

Female heterozygous Crh-iCre mice received bilateral injections (100 nl) of AAV9-CAG-FLEX-ChRmine-EYFP into the ADP (AP = +0.35 mm, ML = ±0.3 mm, DV = –4.9 mm). Control mice received identical injections of AAV9-CAG-FLEX-hrGFP. Following viral injection, dual fibre-optic cannulas (Doric Lenses) were implanted to target either the ADP soma (AP = +0.25 mm, ML = ±0.3 mm, DV = –4.55 mm) or specific downstream projection terminals: the LPO (AP = – 0.45, ML = ±0.8 mm, DV = –5.1), SuM (AP = –2.9 mm, ML = ±0.3 mm, DV = –4.6), or LH/DMH (AP = –1.75 mm, ML = ±0.8 mm, DV = –4.8). The pitches of the dual-fibre cannulas were 0.6 mm for the ADP and SuM, and 1.6 mm for the LH/DMH and LPO.

After 3 weeks of recovery from the surgery, mice were connected to a branching patch cord (200-μm core, 0.37 NA, Doric Lenses) and habituated to the tethering for at least 1 week. Photostimulation was delivered using a 530-nm LED light source (Thorlabs, M530F3) controlled by an LED driver (Doric Lenses, LEDD_2). Light pulses (10-ms duration, 10 Hz, 2-sec on/2-sec off) were applied for 60 min between ZT3 and ZT4. The light intensity at the fibre tip was adjusted to approximately 30 μW for somatic stimulation and 150 μW for terminal stimulation. Surface temperature was continuously monitored at 1-min intervals for 4 h from ZT1 to ZT5 using an infrared thermal imaging camera (Optris PI450) and was defined as the maximum temperature in the thermal image. The recording session comprised a 1-h baseline period, a 1-h pre-stimulation, a 1-h photostimulation period, and a 1-h post-stimulation period. Simultaneously, the thermal camera feed was captured as video at 5 Hz (without temperature metadata) for locomotor analysis. Locomotor activity was quantified using Bonsai software by tracking the displacement of the mouse’s centroid, and the total distance travelled was calculated as an index of locomotion. Although the two data streams were acquired independently, the sampling rates were sufficient to correlate thermogenic responses with physical activity across the 4 h recording period.

### Time-course analysis of BAT thermogenesis and locomotor activity

Female heterozygous Crh-iCre mice received bilateral injections of AAV9-CAG-FLEX-ChRmine-EYFP, were implanted with dual fibre-optic cannulas to target the ADP soma, and were habituated to the tethering as described above. On the first day of habituation, the hair over the interscapular region was shaved under isoflurane anaesthesia to enable reliable estimation of BAT temperature from thermal images. BAT temperature was defined as the maximum temperature in the thermal image. Photostimulation (530nm, 10-ms duration, 5 Hz, 2-sec on/2-sec off) was applied for 10 min, with five trials per mouse performed at 1-h intervals during the light phase. The BAT temperature was continuously monitored at 1 Hz using an infrared thermal imaging camera (Optris, PI450). Locomotor activity was quantified using custom-written Python code by tracking the displacement of the centroid, and the total distance travelled was calculated as an index of locomotion. For ΔT_BAT_ quantification, baseline temperature was defined as the mean temperature during the 5-min period from 10 to 5 min before the onset of each stimulation epoch. The no-stimulation period was defined as the 10-min window beginning 20 min after the end of each stimulation epoch.

### Closed-loop optogenetic stimulation

To investigate the capacity of ADP^Crh^ neurons to actively drive rewarming, we developed a custom closed-loop optogenetic stimulation system based on real-time surface temperature. Mice were fasted at 16 °C as described above. The surface temperature was acquired every 60 s with a thermal imaging camera (Optris, PI450) and analysed in real time using a custom Python script. The closed-loop algorithm was programmed to automatically trigger photostimulation when the surface temperature fell below 33 °C. Upon crossing this threshold, the Python script sent a signal to an Arduino Uno R3 microcontroller, which subsequently delivered TTL trigger pulses to the LED driver. This activated a 530-nm LED (driven at 3 mA) to deliver continuous photostimulation (10-ms duration, 10 Hz, 2-sec on/2-sec off). The system continuously monitored the surface temperature and automatically terminated photostimulation once the temperature rose above 33 °C. Control mice expressing hrGFP underwent the exact same closed-loop protocol.

### Ablation of ADP neurons

Female heterozygous Crh-iCre mice received bilateral stereotaxic injections of AAV2-FLEX-taCasp3-TEVp (100 nl per side) into the ADP (AP = +0.35 mm, ML = ±0.3 mm, DV = –4.9 mm). Control mice received bilateral injections of AAV2-EF1α-DIO-EYFP at the same coordinates and volume. Two weeks after viral injection, mice were implanted intraperitoneally with a temperature logger (DST nano-T, Star-Oddi) under isoflurane anaesthesia. Following at least 14 days of recovery, mice were subjected to torpor, LPS-induced fever, cold-exposure, or cage-exchange stress experiments.

After completion of the behavioural experiments, mice received an intracerebroventricular injection of colchicine (30 µg in 3 µl saline) into the lateral ventricle to enhance somatic accumulation of CRF peptide. Forty-eight hours later, mice were deeply anaesthetized and transcardially perfused, and brains were collected and processed for CRF immunohistochemistry. The extent of ablation was assessed by quantifying the number of CRF-immunoreactive neurons in anatomically matched coronal sections spanning the ADP in control and ablated mice. Mice showing postoperative hemiparesis (*n* = 2) or signs of postoperative inflammation (*n* = 2) were excluded from the analysis.

### LPS-induced fever

Individually housed mice that had received bilateral injections of either AAV2-FLEX-taCasp3-TEVp or AAV2-EF1α-DIO-EYFP into the ADP were injected intraperitoneally with lipopolysaccharide (LPS; 50 µg kg ¹; Escherichia coli O55, Wako, 128-05171) at ZT = 7. Food and water were available ad libitum throughout the experiment. Core body temperature was continuously recorded using an intraperitoneally implanted temperature logger (Star-Oddi, DST nano-T) and analysed using Mercury software (Star-Oddi).

### Cold exposure

Individually housed mice bilaterally injected with either AAV2-FLEX-taCasp3-TEVp or AAV2-EF1α-DIO-EYFP into the ADP were transferred to a temperature-controlled chamber (SHIN-FACTORY, HC-100) maintained at 4 °C at the onset of the dark phase. Food and water were available ad libitum throughout the experiment. Core body temperature was continuously recorded using an intraperitoneally implanted temperature logger (Star-Oddi, DST nano-T) and analysed using Mercury software (Star-Oddi). After 12 h of cold exposure, mice were returned to standard housing conditions.

### Cage-change stress

Individually housed mice bilaterally injected with either AAV2-FLEX-taCasp3-TEVp or AAV2-EF1α-DIO-EYFP into the ADP were transferred to clean cages containing fresh bedding at ZT = 8. Food and water were available ad libitum throughout the experiment. Core body temperature was continuously recorded using an intraperitoneally implanted temperature logger (Star-Oddi, DST nano-T) and analysed using Mercury software (Star-Oddi).

### Fibre photometry

Female heterozygous Crh-iCre mice received a unilateral stereotaxic injection of AAV9-CAG-FLEX-jGCaMP8s (100 nl) into the ADP (AP = +0.35 mm, ML = +0.3 mm, DV = –4.9 mm). During the same surgery, an optical fibre (200-μm core, 0.37 NA, Doric Lenses) was implanted 0.2 mm above the injection site. Two weeks after viral injection, mice were implanted intraperitoneally with a temperature logger (Star-Oddi, DST nano-T) under isoflurane anaesthesia. Following at least 14 days of recovery, individually housed mice were connected to the fibre photometry system via an optical patch cord. GCaMP8s fluorescence signals were recorded using excitation light at 470 nm modulated at 530.481 Hz and 405 nm modulated at 208.616 Hz. The light power measured at the tip of the patch cord was 5-10 μW. Emitted fluorescence was collected through an integrated Fluorescence Mini Cube (iFMC4_AE(405)_E(460–490)_F(500–550)_S; Doric Lenses) and acquired using Doric Neuroscience Studio software (Doric Lenses). The 470-nm channel was used to detect calcium-dependent GCaMP8s fluorescence, whereas the 405-nm channel served as a calcium-independent reference signal. Signals were acquired at 12 kHz, demodulated by the Doric system, and downsampled to 120 Hz for further analysis.

### Data processing and analysis for fibre photometry

Data processing was performed using custom-written Python code. Large-amplitude electrical noise artifacts were removed using a despiking procedure that detected rapid local transients and residual high-amplitude outliers and corrected them based on a median-filtered trace. To improve the signal-to-noise ratio, the despiked signals were low-pass filtered with a Butterworth filter (5 Hz cutoff). To correct for photobleaching, the filtered 405-nm and 470-nm signals were independently baseline-estimated using adaptive iteratively reweighted penalized least squares (airPLS), and the estimated baselines were subtracted to obtain baseline-corrected traces. To remove movement-related artifacts, the baseline-corrected 470-nm signal was regressed against the baseline-corrected 405-nm signal using least-squares linear regression, and the fitted 405-nm component was subtracted from the baseline-corrected 470-nm signal to obtain a residual trace. The residual trace was then divided by the fitted 405-nm signal to calculate ΔF/F, and the resulting ΔF/F trace was z-scored across the recording.

Because the z-scored ΔF/F trace retained temperature-dependent signal changes, calcium events were detected using a local z-score calculated within a 5-min sliding window. Peaks were identified as local z-score events exceeding 3 standard deviations (s.d.). For each detected event, peak magnitude was quantified as the area under the curve (AUC) calculated from the z-scored signal within the full width at half maximum (FWHM) of that event.

For torpor stage classification, body temperature data were linearly interpolated to a 1-min resolution. Torpor entry was defined as a cumulative decrease in body temperature of more than 0.5 °C over the following 10 min. Consecutive time points meeting this criterion were classified as the entry phase. The period from the end of the entry phase until body temperature recovered to within 0.5 °C of the temperature at entry onset was defined as the rewarming phase. One mouse lacking detectable GCaMP expression was excluded from the analysis.

### Whole-brain clearing and light-sheet imaging of ADP^Crh^ neuron projections

Female heterozygous Crh-iCre mice received a unilateral stereotaxic injection of AAV2-CAG-FLEX-synaptophysin–EGFP (50 nl) into the ADP (AP = +0.35 mm, ML = +0.3 mm, DV = –4.9 mm). Three weeks after viral injection, mice were deeply anaesthetized, transcardially perfused with heparin-containing PBS and then 4% PFA, and brains were collected. Brain tissue clearing was performed using an organic solvent-based clearing method, which was based on uDISCO^27^ and FDISCO^28^ methods and empirically optimized to preserve endogenous fluorescence in whole mouse brains. In brief, the brains were immersed in a delipidation solution containing 10% (v/v) 1,2-hexanediol (Tokyo Chemical Industry, H0688), 5% (v/v) Triton X-100, 10 mM N-butyldiethanolamine (Tokyo Chemical Industry, B0725), and 1 mM EDTA at 37 °C for 3 days. After washing three times with PBS at room temperature for 30 min each, the tissues were serially dehydrated with 60% (v/v) tert-butyl alcohol (tBuOH)(KANTO CHEMICAL, 04356-70) at 4 °C for 1 day, 80% (v/v) tBuOH at 4 °C for at least 3 days and 60% (v/v) tBuOH and 40% (v/v) tetrahydrofuran (Wako, 205-17901) at –30°C for 1.5 days. Finally, the samples were cleared in refractive index (RI)-matching solution (RI = 1.554) composed of 50% (v/v) methyl benzoate (Tokyo Chemical Industry, B0074) and 50% (v/v) 2-methylbenzophenone (Tokyo Chemical Industry, M0903) at 4 °C for at least 1.5 days. Dehydration and clearing solutions contained 0.3% (v/v) N-butyldiethanolamine. During all incubation steps, samples were gently shaken. Cleared samples were immersed in an oil mixture (RI = 1.562) composed of silicon oil HIVAC-F4 (Shin-Etsu Chemical, HIVAC-F4) and HIVAC-F5 (Shin-Etsu Chemical, HIVAC-F5), and imaged using a light-sheet fluorescence microscope, UltraMicroscope Blaze (Miltenyi Biotec). Images were acquired using a 1.1× objective, NA 0.1, MI PLAN lens with a 2.5x optical zoom.

### Anterograde tracing

Female heterozygous Crh-iCre mice received a unilateral stereotaxic injection of AAV2-CAG-FLEX-synaptophysin–EGFP (50 nl) into the ADP (AP = +0.35 mm, ML = ±0.3 mm, DV = –4.9 mm). Three weeks after viral injection, mice were deeply anaesthetized, transcardially perfused with PBS and then 4% PFA. The brains were collected, post-fixed in 4% PFA solution at 4 °C overnight and immersed in 30% sucrose PBS at 4 °C for at least 2 days. A series of 60-µm-thick sections was obtained using a Leica CM3050S cryostat (Leica Microsystems) or RWD Minux FS800A cryostat. EGFP signal was amplified by immunohistochemistry using an antibody against GFP. Images were obtained using a BZ-9000 fluorescence microscope (Keyence).

### Slice preparations and Whole-cell recordings

For slice preparation, mice were deeply anaesthetized via intraperitoneal (i.p.) injection with a mixed anesthetic solution containing 40% ketamine hydrochloride (Ketalar® injection 500 mg, Daiichi Sankyo Co., Ltd.) and 15% xylazine hydrochloride (Selactar® 2% injection, Elanco Japan K.K.), diluted in saline. Mice were intracardially perfused with ice-cold cutting solution containing (in mM): 93 NMDG, 2.5 KCl, 1.2 NaH_2_PO_4_, 30 NaHCO_3_, 20 HEPES, 25 glucose, 5 sodium ascorbate, 3 sodium pyruvate, 0.5 CaCl_2_, 10 MgSO_4_, and 12 N-acetyl-L-cysteine (pH 7.3), bubbled with 95% O_2_ and 5% CO_2_. Coronal brain slices (300 µm thick) containing the LPO were prepared using a Neo Linear Slicer MT (D.S.K, Osaka, Japan). The slices were transferred to warmed (30 °C) cutting solution for 10 min for recovery, then were kept in oxygenated artificial cerebrospinal fluid (aSCF) containing (in mM): 125 NaCl, 2.5 KCl, 1.25 NaH_2_PO_4_, 26 NaHCO_3_, 20 glucose, 1 MgSO_4_, and 2 CaCl_2_ and subsequently maintained at room temperature until use.

Whole-cell voltage-clamp recordings were performed from LPO neurons using an IPA amplifier (Sutter Instruments). Signals were sampled at 20 kHz and acquired with SutterPatch software (Sutter Instruments). Recordings were conducted at room temperature using a differential interference contrast microscopy (Evident Corporation, BX51WI) equipped with an IR-CCD camera (IR1000, DAGE-MTI) and an LED light source (470-nm; CoolLED Ltd.). Slices were continuously perfused with oxygenated ACSF.

Patch electrodes (2–4 MΩ) were filled with an internal solution containing (in mM): 140 K-gluconate, 20 KCl, 2 MgCl_2_, 0.2 EGTA, 10 HEPES, 2 Mg-ATP, 0.5 Na-GTP (pH 7.4, adjusted with KOH, 290 mOsm). Series resistance was not compensated. Full-field optical stimulation of the maximum light intensity (59.2 mW/mm²) (100%) with 300 µs duration was delivered every 60 sec using 470-nm light through a 40x objective to evoke ChR2-mediated synaptic currents. To pharmacologically isolate light-induced monosynaptic responses, the following antagonists were bath-applied: 0.5 µM tetrodotoxin (voltage-gated Na^+^ channel blocker; Fujifilm Wako Pure Chemical Corporation); 200 µM 4-aminopyridine (4-AP) (voltage-gated K^+^ channel blocker; Kanto Chemical Co. Inc.); 10 µM NBQX (AMPA receptor antagonist; Tocris Bioscience); 10 µM R-CPP (NMDA receptor antagonist; Tocris Bioscience); 10 µM bicuculline methochloride (GABA_A_ receptor antagonist; Tocris Bioscience). All compounds were prepared as stock solutions according to the manufacturer’s instructions and diluted to their final concentrations in ACSF immediately before use. The amplitude of light-induced synaptic responses after application of glutamate receptor or GABA receptor antagonists was calculated as a percentage of the synaptic response amplitude measured after application of TTX and 4-AP.

### Statistics

All data were presented as the mean ± standard error of the mean (s.e.m.). Statistical analyses were performed using GraphPad Prism version 9.0.2 for macOS (GraphPad, USA). A *p*-value less than 0.05 was considered statistically significant. To test normality, the Shapiro–Wilk test was performed.

## Acknowledgments

We thank members of the Yamanaka, Ono, and Wake laboratories for discussions; members of the Ono laboratory for reagents; R. Nozaki, S. Tsukamoto, T. Kobayashi, and S. Nozaki for technical assistance. We thank the core facility of the Nagoya University Graduate School of Medicine for the use of the Ultramicroscope II and S. Kato at the Brain/MINDS 2.0 vector core for virus packaging. This work was supported by JSPS KAKENHI grants to H.Y. (23K26830, 24H02007, and 25K22336), by JST PRESTO to H.Y. (JPMJPR21SA), and by the Takeda Science Foundation, the Lotte Foundation, and the Hori Sciences and Arts Foundation to H.Y. A.O. was supported by JSPS KAKENHI (23H04939), Takeda Science Foundation scholarship and Nagoya University CIBoG WISE program of the Ministry of Education, Culture, Sports, Science and Technology (MEXT).

## Contributions

A.O. and H.Y. conceptualized the work. A.O. designed, performed and analysed the experiments. A.O. and N.F. analysed the whole-brain immunostaining data. M.N. performed electrophysiological recordings in brain slices. A.O. and C.J.H. performed in situ hybridization experiments under the supervision of D.O. U. A. collected mouse brain samples during torpor rewarming. C.S. collected hamster brain samples under the supervision of Y.Yam. H.U. and K.T. performed brain tissue clearing and light-sheet fluorescence imaging. A.Y., S. T.-K. and H.W. contributed reagents and advised on the study. H.Y. supervised the study. A.O. and H.Y. wrote the manuscript. H.Y. acquired funding.

